# Brainprints: identifying individuals from magnetoencephalograms

**DOI:** 10.1101/2020.06.18.159913

**Authors:** Shenghao Wu, Aaditya Ramdas, Leila Wehbe

## Abstract

Magnetoencephalography (MEG) is used to study a wide variety of cognitive processes. Increasingly, researchers are adopting principles of open science and releasing their MEG data. While essential for reproducibility, sharing MEG data has unforeseen privacy risks. Individual differences may make a participant identifiable from their anonymized recordings. However, our ability to identify individuals based on these individual differences has not yet been assessed. Here, we propose interpretable MEG features to characterize individual difference. We term these features brainprints (brain fingerprints). We show through several datasets that brainprints accurately identify individuals across days, tasks, and even between MEG and Electroencephalography (EEG). Furthermore, we identify consistent brainprint components that are important for identification. We study the dependence of identifiability on the amount of data available. We also relate identifiability to the level of preprocessing, the experimental task. Our findings reveal specific aspects of individual variability in MEG. They also raise concerns about unregulated sharing of brain data, even if anonymized.

Figure 1:
Graphical abstract.
Identifying which subject a segment of MEG data belongs to is strikingly easy when other data from the same session is available for every subject. We propose three types of interpretable features that can also be used to identify individuals across sessions with high accuracy. Identifiability of individuals is influenced by factors such as resting state vs. task state, components of each feature, the sample size and the level of preprocessing. Our results reveal aspects of individual variability in MEG signals and highlight privacy risks associated with MEG data sharing.

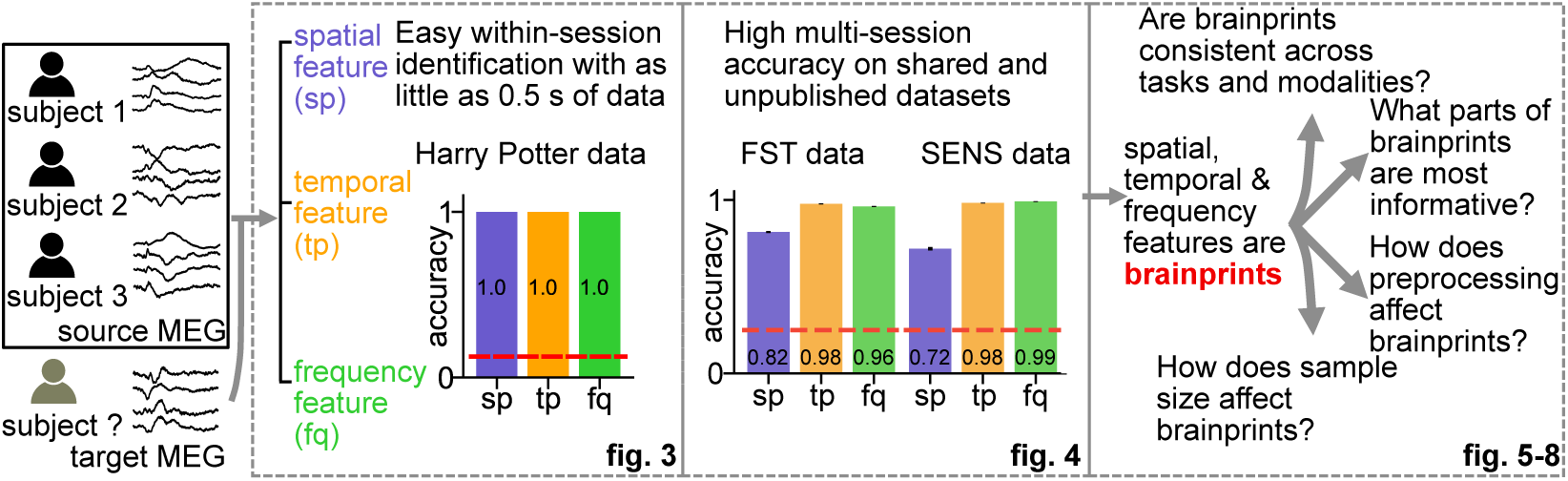

## Introduction

The open science movement is a result of the increasing awareness of the importance of sharing data and code to promote scientific reproducibility [1]. Public repositories enable researchers to share their neuroimaging data (fMRI, EEG, MEG, etc.) while making sure to censor out individual information [2]. However, data anonymization does not always preserve privacy [3]. Combining different types of information using methods such as record linkage approaches [4] may cause serious privacy violations. This problem is exacerbated when multiple datasets that happen to contain the same individual are available, which is rather common in neuroimaging (e.g. [5]). Hence it is natural to ask if anonymized individuals can be identified from neuroimaging datasets and if so to what degree. Specifically, we ask: do individuals have a *brainprint*, a brain-activity analog of a fingerprint? If there is evidence for a brainprint, then researchers may be warned about how easily individual information can be inferred, and it may cause them (and the field) to act with more caution when publishing neuroimaging data online. For instance, it may pave the way for the adoption of more sophisticated data-release mechanisms like differential privacy [6] and homomorphic encryption [7].

Assume there are two multi-individual neuroimaging datasets with overlapping participants: a “source” dataset and a “target” dataset. The question of interest is: can we accurately decide which individual in the source dataset corresponds to the individual in the target set? In other words, is there individual *identifiability* between the two datasets? The aforementioned question could arise naturally in practice: it is very common for university labs to recruit their own lab members for preliminary studies; these are anonymously released with an associated publication. Assume that one year later, lab member A relocates to city B, and privately volunteers for a study by a public hospital that tracks the effect of a drug (or some intervention) on patients in early stages of early-onset Alzheimer’s, while collecting MEG data. If this data is also anonymously released at a future point, brainprints could plausibly be used to detect a common participant, thus identifying that A has Alzheimer’s because only one member of the lab moved to city B. This would already be a gross unintended violation of privacy, but one can further imagine that an insurance company uses this to prove that a condition was pre-existing at the time of the first scan (before the individual themselves knew), or use it to decide individual-level pricing.

If high individual identifiability exists even if the source and target set were recorded in separate sessions for each individual, there might be essential differences in the patterns of the data among individuals which is preserved across scanning sessions. Namely, individual identifiability might be related to variability in brain structure or function (or other individual characteristics such as head size). In multi-individual, multi-session neuroimaging data, there exists “within-session” variability across individuals in the same session and “cross-session” variability of the same individual cross sessions [8]. For simplicity, consider the four scenarios in Fig. 2. Low variability in both within-session (individuals are similar) and cross-session (an individual’s data is consistent across session) is likely to promote statistical power for detecting average group effects with fixed sample size, thereby facilitating reproducibility [9, 10]. High cross-session and low within-session variability (e.g. individual 1’s data in session 1 is very different from their data for session 2, but somehow very similar to individual 2’s data in session 1) may indicate session-specific artifacts (e.g. the scanner was faulty during the recording of session 1 for all individuals). High cross- and within-session variability makes data unreliable. Finally, high within-session (individuals are different from each other) and low cross-session variability (individuals are similar to themselves) leads to individual identifiability. Individual identifiability in turn indicates *consistent* individual differences, which in themselves are an important topic of scientific enquiry [8, 11]. Understanding sources of consistent variability can help learn the underpinnings of disease or more generally to map the relationship of brain structure and activity to individual behavioral characteristics.

**Figure 2:**
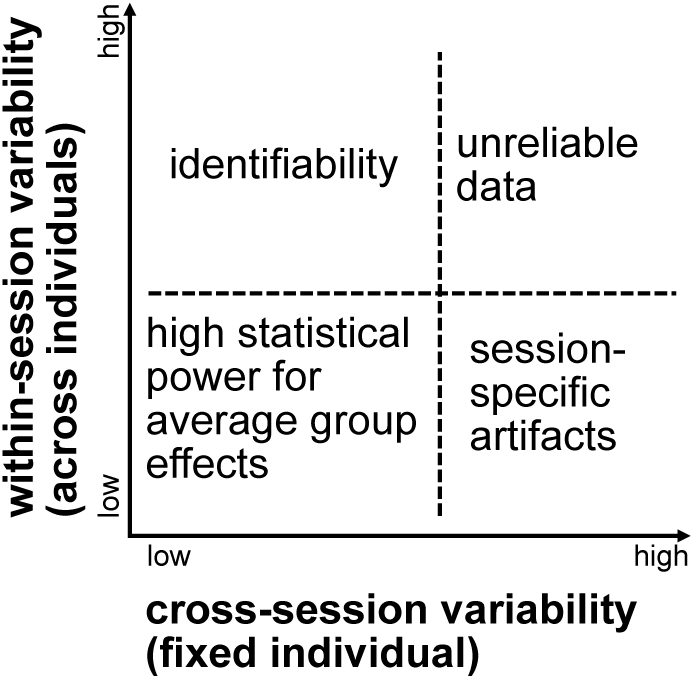
Individual identifiability is a function of individual and session variability in neuroimaging. Consider repeating an experiment in multiple sessions for a group of individuals. Cross-session variability refers to the change in the recorded data for the same individual across sessions, while within-session variability refers to differences in a single session’s recorded data across individuals (keeping all other variables, including the stimulus, unchanged). The ideal conditions for the scientific discovery of an effect shared by the group is low within-session and low cross-session variability. However, the combination of low within-session and high cross-session variability indicates an artifact or a confound in the experiment design (e.g. each month, one session is recorded for all individuals and the instrument has a drift over time). High within-session variability paired with low cross-session variability leads to individual identifiability with the individual’s data acting like a stable signature that differentiates them from others. Finally, high within-session and cross-session variability lead to unreliable data.

Similar individual identification problems have been studied using EEG and fMRI for the purpose of biometric authentication and to investigate individual differences [12, 13, 14, 15, 16, 17, 18]. The term ‘brainprint’ has also been previously used to represent brain-specific information, such as morphology and event related potential biometrics [19, 20, 21], that can be used to identify individuals. Individual identification with MEG data, however, has not been fully explored. Due to availability of MEG datasets, only single-trial MEG data has been studied for person identification [22]. Other MEG studies focusing on variability of individual data [8, 11] may not make connections with individual identifiability. Cross-modality identifiability has also not been explored.

In this paper, we argue that individuals can be easily identified with MEG data. We measure identifiability as identification accuracy with three interpretable MEG features on multiple public and private MEG datasets. We show that identifiability is not a product of environmental artifacts and specific features have a consistent performance between task and resting state data. We further demonstrate that identifiability is preserved even between MEG and EEG datasets. We dissect the contribution of each features into sub-features to understand what may be leading to the high identifiability. Factors such as the amount of data and level of preprocessing are also shown to have influence on identifiability. Our analysis not only confirms the worrisome potential of privacy being compromised by released MEG data via extracting simple features but also leverage the interpretability of the features to explain the underlying mechanism for the high identifiability, thereby relating it to individual variability.

## Results

In the result section, we first confirm the existence of the fingerprint using data of a single session, of multiple-sessions, of multiple-tasks, and even of multiple recording modalities. We use machine learning tools as well as interpretable features to show that identification is easy when the MEG sessions were collected on a single visit. We then show that the proposed features also achieve high accuracy on datasets of multiple visits to the scanner, and some features are even consistent on datasets between different tasks and from imaging recording modalities. We finally show which components of each feature is important for individual identification, and that sample size and level of preprocessing will also affect identification accuracy.

### Within-session identification is surprisingly easy

To measure identifiability, we consider the test accuracy of a classifier trained to identify participants from their MEG recording. We first focus on within session identifiability. In this context, we assume that each participant undergoes one session. A classifier is trained on a subset of the session, in which each trial is labeled with the identity of the participant it corresponds to. In our framework, we refer to the training set as the source set. Then, on held-out test data, the classifier predicts which participant is associated with each test trial. We refer to the test set as the target set. As an example, we investigated individual identifiability on a MEG dataset of *eight* participants during a reading task. Participants were asked to read a chapter of Harry Potter [23] while each word was presented for 0.5 s on a screen. There were 5176 trials (words) for each individual. The data was recorded using the Elekta Neuromag system. The Harry Potter (HP) data is a single-session dataset: the data for each individual were collected on a single visit of the MEG scanner. Hence the source and target set are non-overlapping subsets of that single session. We preprocessed and downsampled the data from 1000 Hz to 200 Hz so that there are 100 time points for each word. We trained a random forest classifier [24] using the MEG recording of all channels at a randomly selected time point, a flattened vector representing the snapshot of the topographic map (topomap) of the brain activity (see Methods and Fig. S3). Random forest is a powerful classifier that uses a majority vote of a number of decision trees to predict the label associated with a given feature. Under this setting, we are asking if there is any individual-specific information contained in the topomap, the basic element of MEG recording. We split the dataset into 10 non-overlapping folds and used one as the target (testing) set and the other nine as the source (training) set. This 10-fold cross-validation scheme yielded a high identification accuracy (0.94) while the chance accuracy is only 0.125. We also repeated the analysis by only sampling one topomap from each trial to deflate possible statistical dependency and still obtained an accuracy of 0.923. This surprisingly high accuracy on merely 0.05 s of MEG data suggests the existence of strong patterns detected by the random forest classifier. This strong pattern may be contained on the transient spatial distribution of an individual’s MEG activity and is strongly distinctive of an individual. This high accuracy with the limited amount of information used suggests that within-session identification is a strikingly easy task.

### Interpretable MEG features yield high identification accuracy

The random forest classifier may not enclose enough information to explain the high identifiability of the HP data because of the black-box nature of the algorithm. The topomap mainly contains the spatial information: how heterogeneous the amplitude of the signal is across channels at a certain time point. High identifiability may also be attained using temporal and frequency information. We proposed three interpretable features for individual identification to further disseminate the individual-specific information. These features are interpretable because they bear biological meanings and hence can be used to interpret the high identification accuracy. The three features were based on *n* randomly selected trials (words) which have the shape [102 channels, 100 time points, *n* trials] (Fig. 3(a)). **sp** (Fig. 3(b)) is the spatial correlation between different sensors which may be related to individual-specific correlated activities between areas of the brain or the anatomy of the individual (e.g. brain size) [8, 25]. **tp** (Fig. 3(c)) is the temporal correlation between the time points into a trial. A high value in the **tp** matrix indicates highly synchronous brain signals between two temporal points, which might be related to participant specific stimulus processing latencies. A relevant study shows that the temporal change of brain activities in auditory steady-state responses are different between individuals [26]. **fq** (Fig. 3(d)) represents the distribution of the power intensity of signal frequency. Individual differences might also manifest as differences in the power distribution along frequency bands [27, 22].

**Figure 3:**
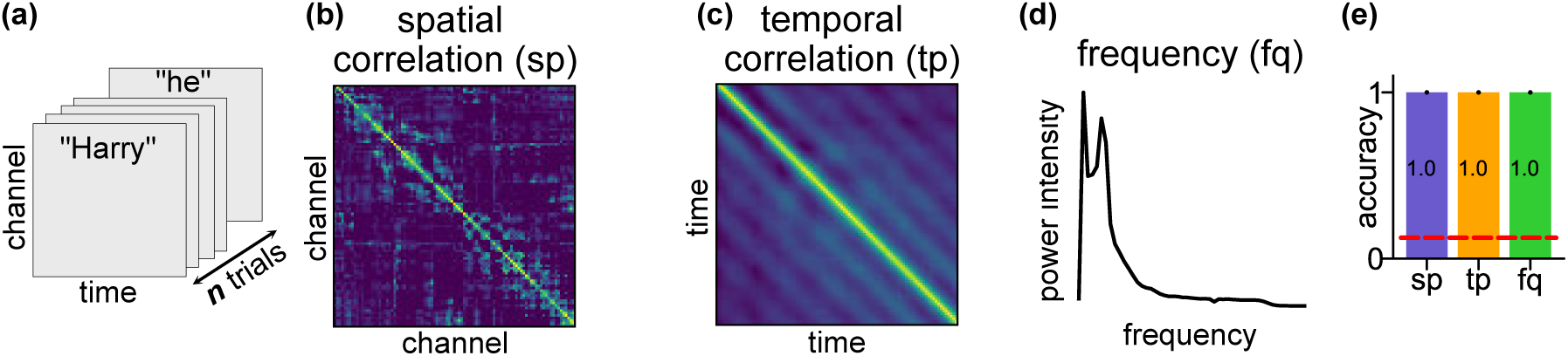
High within-session identification accuracy on HP data with three interpretable features. **(a)** Shape of the HP data before featurization. The HP data consists of participants reading a book chapter one word at a time for 0.5s each. The data is resampled to have the dimension [102 channels, 100 time points, *n* trials] where each trial corresponds to one word and *n* to the number of words. **(b)** The spatial correlation feature **sp** is a 102 × 102 Pearson’s correlation coefficient matrix computed across the time points and trials. **(c)** The temporal correlation feature **tp** is a 100 × 100 Pearson’s correlation matrix computed across the channels and trials. **(d)** The frequency feature **fq** is a vector in ℝ^51^ where 51 is the number of frequency bands. The power at each band was averaged across channels and trials. **(e)** Identification accuracy with the three features. The accuracy was averaged across 100 identification runs of 8 individuals. The red dashed line represents the chance level (= 0.125). The error bars are the standard errors across individuals and identification runs and are invisible since they are all zeros.

We used the 1-Nearest Neighbor (1NN) identification procedure, similar to Finn et al. [15], to test if the three features are *brainprints* for the within-session identification task. For a given feature such as **sp**, the feature was computed on the source set using *n* randomly sampled trials (*n* = 300 for the HP data). Target set features were also computed in the same way (but unlabeled) with the same number of trials. The 1NN classifier simply assigned each target feature to the participant with the closest source feature (we used correlation to measure distance). The aforementioned matching process was repeated for 100 runs to account for the variance of the feature on the sampled trials, and the accuracy was averaged across these 100 runs. The simplicity of this 1NN classifier optimizes the interpretability of the result. With *n* = 300 trials all three features achieve perfect identification accuracy (Fig. 3(e), the accuracy for **sp, tp**, and **fq** is 1 ± 0, mean ± SE, *p <* 0.0002, see Supplement B for how we computed the p-values). In fact, the high identifiability can be attained with as few as *n* = 100 trials (Fig. S2(a)). The high identifiability with **sp, tp** and **fq** suggests they are *brainprints*, at least for identifying individuals within a session. Therefore, multiple features capturing different aspects of the MEG activity can be used for identifying individuals.

### Cross-session identification confirms the existence of brainprints

The high within-session identification accuracy suggests **sp, tp**, and **fq** are individual-specific within a session. Artifacts such as environmental noise and equipment configurations, however, might be the main contributing factor to within-session identification accuracy. Hence, we examined the consistency of the three features when the same type of task data was collected from each individual on multiple sessions. This setting tests if the features are preserved over time, i.e. if they are indeed *brainprints* and not mere artifacts. If the identifiability is significantly lower on multi-session datasets, the high identifiability on the HP data may be a mere result of session-specific artifacts, since the recording session for each individual is performed on different days. If high cross-session identifiability is observed, **sp, tp**, and **fq** can be considered genuine brainprints because they are unique to individual and invariant between sessions. This would also suggest low cross-session and high within-session variability (Fig. 2).

We tested the three features on two multi-session datasets: FST [28], a four-session dataset where four individuals were shown pictures of familiar and unfamiliar faces with 1464 trials and SEN, a three-session dataset where four individuals were shown sentences with 3575 trials. Both recordings were recorded with the Elekta Neuromag sytem and were preprocessed and downsampled from 1000 Hz to 200 Hz so that there were 100 time points in one picture/sentence which we considered as one trial (see Methods). Since each individual has recordings conducted on different days, we set the target and source data to be from different sessions (Fig. 4(a)), to test the role of environmental artifacts and further confirm the existence of the brainprints. In addition to identification accuracy which is binary on one matching procedure, we used a continuous version, the *rank accuracy*, which captures more information in a failure case where an individual is misidentified. Rank accuracy captures the rank of the correct assignment out of all possible assignments; it is 1 if the target feature of each individual have the largest similarity to the source features for that individual, and is 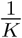 if the similarity is the smallest. The chance rank accuracy is 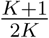. In addition to the identification and rank accuracy, we also used a metric, differential identifiability [29] which measures the similarity between the features of the same individual as compared to that of other individuals (see Methods and Fig. S9).

**Figure 4:**
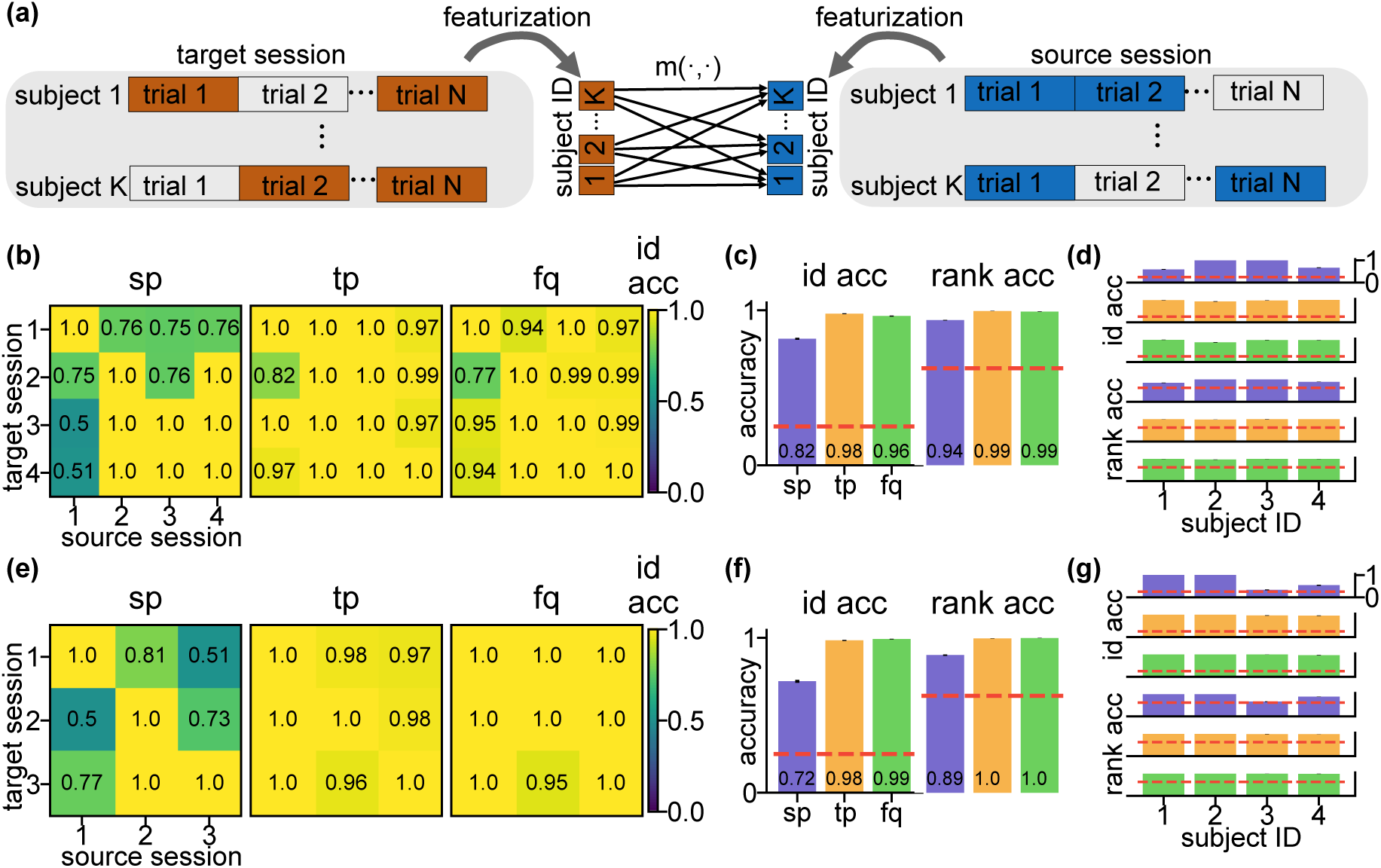
Cross-session identification on FST and SEN data confirms existence of brainprints. **(a)** Schema of the cross-session identification task. For one identification run, the features of each individual are computed using randomly sampled trials (*N* = 300) from both the source and target session. Target session features are then classified by selecting the individual with the largest similarity score in the source session. **(b)** Heat maps of the cross-session identification accuracy using the three features on FST data. Each grid represents the average accuracy across 4 individuals and 100 identification runs. The within-session accuracy (diagonal entries) are computed using the same source-target splitting procedure as on the Harry Potter data to avoid data leakage. **(c)** Average cross-session identification accuracy and rank accuracy for each feature on FST data. Within-session accuracy (diagonal entries in **(b)**) were excluded in computation. Error bars are the SEs across cross-sessions (*N* = 12), individuals (*N* = 4), and identification runs (*N* = 100) and are invisible due to small values. Red dashed lines are the chance level for the identification accuracy (= 0.25) and rank accuracy (= 0.625). **(d)** Identification and rank accuracy on FST data by individual. Within-session accuracy were excluded in computation. Error bars are the SEs across cross-sessions (*N* = 12) and and identification runs (*N* = 100) and are invisible due to small values. The red dashed lines are the same as in **(c). (e)–(g)**, same as **(b)–(d)** but on SEN data with the same number of individuals and identification runs (*N* = 4 and *N* = 100) but different number of cross-sessions (*N* = 6). The high identification accuracy with the three features on multi-session datasets confirms these features can be brainprints for individual identification.

Both **tp** and **fq** achieve almost perfect average identification and rank accuracy on both FST and SEN data whereas **sp** achieved lower but still well above-chance accuracy (Fig. 4(c,f)). The high cross-session identification accuracy of **sp, tp**, and **fq** confirms that it is reasonable to call them brainprints for individual identification in MEG. The lower identification accuracy for **sp** is due to low accuracy on a two of the individuals (Fig. 4(d,g)) in both datasets. This is also confirmed using the confusion matrices (Fig. S7 However, identification accuracy of these individuals is not consistently low across all session pairs (Fig. 4(b,e)) indicating that **sp** only perform worse for these subjects between certain sessions.

For SEN data, the MEG recording of two subjects were taken on the same day for session 1 and 2. Since the identification accuracy of **sp** corresponding to these two pair of sessions (1 vs 2 and 2 vs 1) does not yield higher accuracy than the average (the mean identification accuracy between these two session pairs is 0.655, lower than 0.72, the mean across all cross-session pairs), the accuracy for **sp** is not inflated due to this issue with duplicated recording times. In line with the results on the HP dataset, **sp, tp**, and **fq** are the brainprints that are consistent even between recording sessions with **tp, fq** leading to higher identifiability.

### Spatial brainprints are consistent across resting-state and tasks

The high performance and interpretability of the brainprints make it enticing to study the factors and the underlying mechanism for identification. We looked at the performance of these features between two sessions of different types collected on the same day to test their consistency between different brain states. We compared the features using the Human Connectome Project (HCP) MEG data [5] between a resting-state session (422 trials in average) in which individuals (*N* = 77) rest and do not perform a task and a task-MEG session (372 trials in average) where these same individuals view images and perform a working memory task. The dataset was recorded using the MAGNES 3600 system. We preprocessed and downsampled the data from 1024 Hz to 200 Hz and there were 500 time points in one trial in the WM data which corresponds to 2.5 s after the onset of the stimulus (see Methods). For the resting dataset, we simply reshape the recording into consecutive blocks similar to the WM dataset and performed the same analysis.

Consistent with the cross-session results in Fig. 4, **sp** yields a high identification accuracy (Fig. 4(b), 0.77 ± 0.0034, mean ± SE, *p <* 0.0002), well above the 0.013 random baseline. This suggests that the spatial fingerprint is consistent between different brain states which confirms a similar finding in fMRI [15]. The by-individual identification accuracy (Fig. 5(c)) shows that there is a small subset of individuals whose accuracy is below random, which may be due to the lack of head position correction in the HCP collection protocol. **tp** and **fq** do not perform as well as **sp**, suggesting that the temporal rhythm and frequency involved might be different between resting-state and task [30, 31].

**Figure 5:**
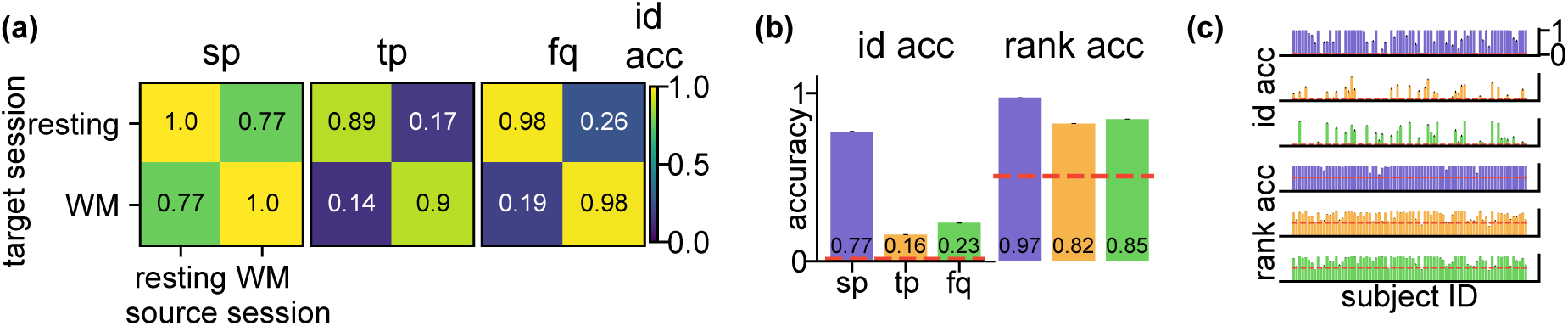
Consistent sp for cross-task identification on Human Connectome Project data. **(a)** Heat maps of the cross-task identification accuracy using the three features on HP data. Both resting and working memory (WM) data were recorded on the same day. For one identification run, the features of each individual were computed using randomly sampled trials (*N* = 200) from both the source and target session. Each grid represents the average accuracy across 77 individuals and 100 identification runs. The within-task accuracy (diagonal entries) were computed using the same source-target splitting procedure as on the Harry Potter data to avoid data leakage. **(b)** Average cross-task identification accuracy and rank accuracy for each feature on HCP data. Within-task accuracy (resting vs. resting, WM vs. WM) are excluded in computation. Error bars are the SEs across cross-task sessions (*N* = 2), individuals (*N* = 77), and identification runs (*N* = 100) and are invisible due to small values. The red dashed lines are the chance level for the identification accuracy 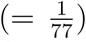 and rank accuracy 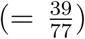. **(c)** Identification (upper three rows) and rank (lower three rows) accuracy on HP data by individual. Within-task accuracy were excluded in computation. Error bars are the SEs across cross-task sessions (*N* = 2) and identification runs (*N* = 100) and are invisible due to small values. The red dashed lines are the same as in **(b)**. These results indicate that **sp** is consistent even when performing different tasks (resting vs WM) in the source and target session.

The rank accuracy of **tp** and **fq** (Fig. 5(b), 0.82 ± 0.0017 and 0.85 ± 0.0016, mean ± SE, *p <* 0.0002 for all) are much higher than the baseline (= 0.506). The majority of the individuals also have higher rank accuracy than baseline for **tp** and **fq** (Fig. 5(c)). The higher rank accuracy suggests that **tp** and **fq** may still contain individual-specific information but are not strong enough to achieve a high identification accuracy. Since the individuals perform different tasks on the source and target session, the rank accuracy indicates the potential consistent brainprint the generalizes beyond the task. It is noticeable that for the HCP dataset, the recording sessions of one individual were recorded on the same day. Hence one may exercise caution when extend the conclusions to cross-session datasets.

### Temporal and frequency brainprints are consistent across modalities

So far we have verified that the brainprints are consistent across visits, and even between resting and tasks. It would be a stronger piece of evidence if we show that brainprints can identify individuals during two visits to different centers with different recording modalities. We looked at MEG and EEG (electroencephalography) data of 14 participants viewing scene images (362 trials for each individual, one trial lasted 1 s), obtained from [32]. Both MEG and EEG were recorded for the exact same stimuli, but on different days for each participant, making it an ideal testbed to verify the consistency of brainprints across different imaging modalities. We downsampled the MEG and EEG data from 1000 Hz and 512 Hz to 110 Hz so that there were 110 time points per trial. Since the spatial arrangements of MEG and EEG are different, we only tested the accuracy using **tp** and **fq**.

Both features yield well above-chance identification accuracy and rank accuracy (Fig. 6, identification accuracy for **tp** and **fq** is 0.45 and 0.46 whereas the chance accuracy is 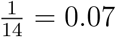, same conclusion for the rank accuracy). This constitutes strong evidence that the frequency and temporal information of an individual’s response to stimuli are preserved even when different imaging modalities are used. The consistency also indicates that, at least for the temporal and frequency feature, the high accuracy is due to the individual-specific responses despite the possibility of different artifacts induced by MEG and EEG machines.

**Figure 6:**
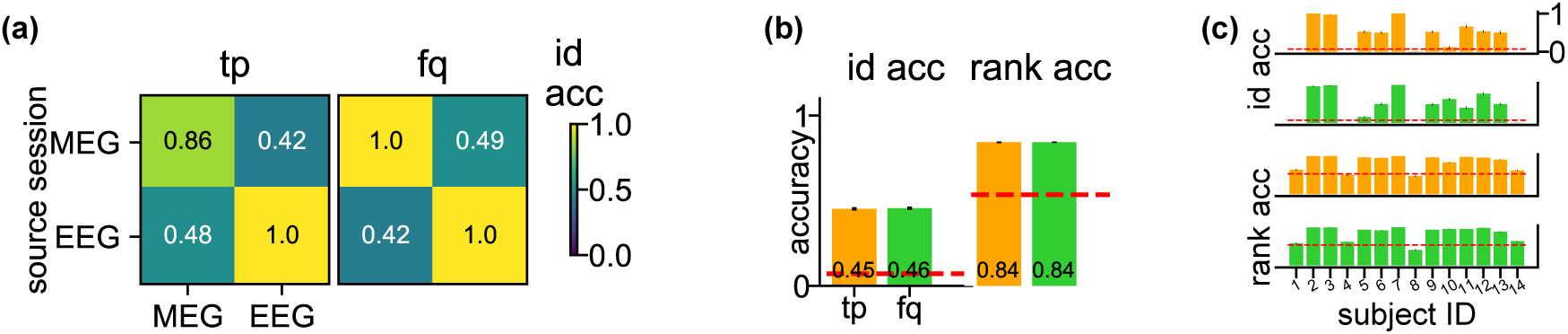
Consistent tp and fq for cross-modality identification on MEG-EEG data. **(a)** Heat maps of the cross-modality identification accuracy using the two features on MEG-EEG data. MEG and EEG data for the same individual were recorded on different days. For one identification run, the features of each individual were computed using randomly sampled trials (*N* = 200) from both the source and target session. Each grid represents the average accuracy across 14 individuals and 100 identification runs. The within-task accuracy (diagonal entries) was computed using the same source-target splitting procedure as on the Harry Potter data to avoid data leakage. **(b)** Average cross-modality identification accuracy and rank accuracy for each feature. Within-modality accuracy (MEG vs. MEG, EEG vs. EEG) were excluded in the computation. Error bars are the SEs across cross-modality sessions (*N* = 2), individuals (*N* = 14), and identification runs (*N* = 100) and are invisible due to small values. The red dashed lines are the chance level for the identification accuracy 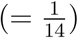 and rank accuracy 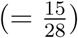. **(c)** Identification (upper two rows) and rank (lower two rows) accuracy on MEG-EEG data by individual. Within-modality accuracy is excluded in the computation. Error bars are the SEs across cross-modality sessions (*N* = 2) and identification runs (*N* = 100) and are invisible due to small values. The red dashed lines are the same as in **(b)**. These results indicate that **tp** and **fq** are consistent even when different neuroimaging modalities were used in the source and target session.

### Not every part of a brainprint is equally important

What contributes to the high identifiability of the three brainprints? Understanding the relative contribution of the components of brainprints could help understand individual identifiability and variability. We divided the three brainprints into sub-features and looked at their identification accuracy to see which components contain the most individual-specific information. **sp** was divided into correlations between groups of sensors: Left Occipital (LO), Right Occipital (RO), Left Parietal (LP), Right Parietal (RP), Left Temporal (LT), Right Temporal (RT), Left Frontal (LF), Right Frontal (RF). **tp** was divided into correlations between time intervals. **fq** was divided into frequencies within a sliding window. We use the SEN and FST dataset to focus on cross-session patterns.

For both SEN and FST, the correlations between sensors within Left Occipital (LO) and between LO and Right Parietal (RP) yielded high accuracy (Fig. 7(a), inset, and Fig. S10). LO is involved in visual processing [33] and RP is involved in sensory integration [34], both of which are functions recruited by the experimental task. Due to the nature of the sampled signal and the physical properties of the skull, each MEG sensor samples coarsely from the brain, making it hard to say whether MEG spatial correlation effectively corresponds to functional connectivity, especially for nearby sensors [8]. However, the fact that correlations between faraway groups of sensors, for example, LT and RT, still have good accuracy suggesting it may be due to actual functional correlation between these areas, but it could still be the case that it is the difference in skull shapes that contributes to the high **sp** accuracy.

**Figure 7:**
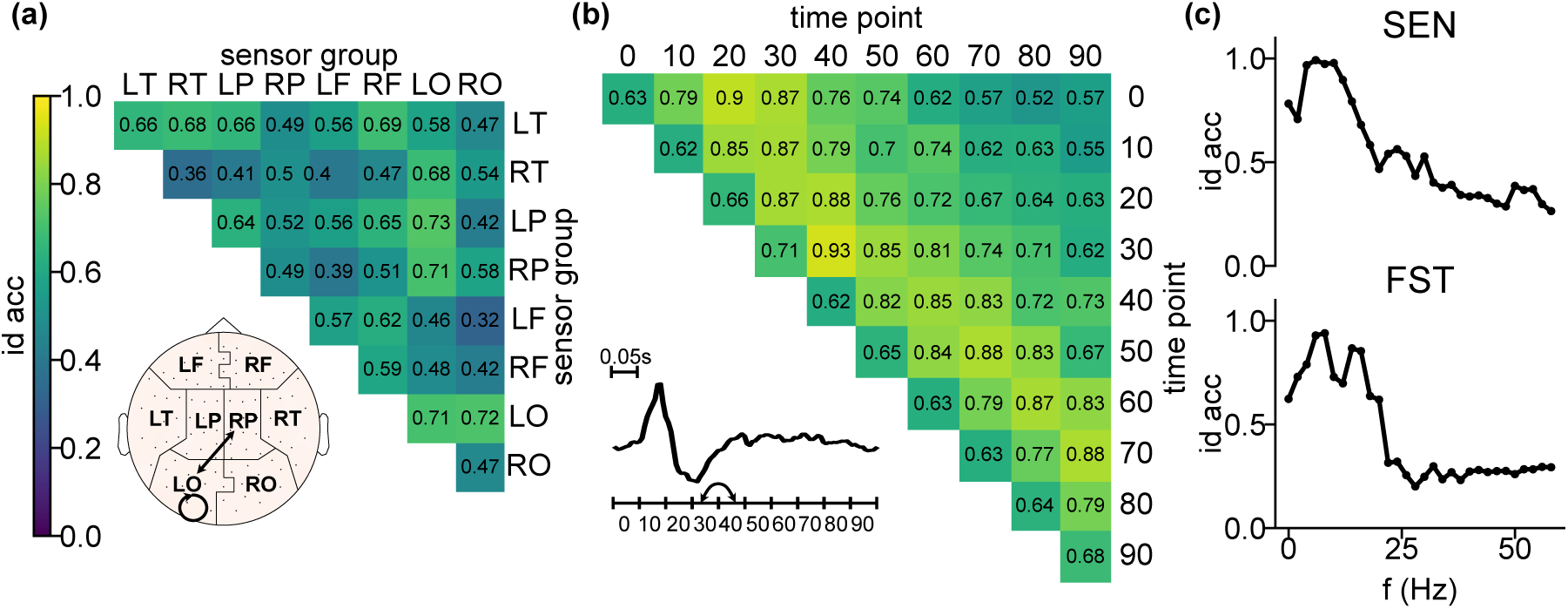
Identification accuracy of components of the features. See Fig. S10 for **a,b** on FST data. **(a)** Identification accuracy of the sub-features of **sp** on SEN data. Each grid represents the identification accuracy using the corresponding entries of **sp** averaged across cross-sessions (*N* = 6), individuals (*N* = 4), and identification runs (*N* = 100). Inset is the plot of the sensor group layout and edges correspond to the sensor group pair with over 0.7 accuracy for both FST and SEN. The topomap was plotted using the python MNE package [37]. **(b)** Identification accuracy of the sub-features of **tp** on SEN data. Each grid represents the identification accuracy using the corresponding entries of **tp** averaged across the same dimensions as in **a**. Inset is an example MEG signal of one individual averaged across channels (*N* = 102) and trials (*N* = 1000). Arrows correspond to the entries of the heatmap with over 0.9 accuracy for both FST and SEN. **(c)** Identification accuracy of the sub-features of **fq** on SEN (upper plot) and FST (lower plot) data. Each dot represents the identification accuracy using the corresponding entries of **tp** averaged across cross-sessions (*N* = 6 for SEN and 12 for FST), individuals (*N* = 4), and identification runs (*N* = 100). Accuracy values of *f* larger than 60 Hz were truncated since the curve became flat. Error bars are SE across cross-sessions, individuals, and identification runs and are invisible due to small values. The curve peaks at *f* = 6 Hz for SEN and *f* = 8Hz for FST. The accuracy of some components of a feature is consistently higher than the rest on both datasets, indicating that some parts of a certain feature may be more important in identifying individuals.

For both SEN and FST, the super-diagonal of the heat map for temporal sub-features (Fig. 7(b) and Fig. S10) had high accuracy. The super-diagonal entries correspond to the cross-correlation of the MEG signal between two consecutive segments of 0.05 s. Hence the rhythm of the signal within a short segment of time contributes to identifiability, which can also be seen from the banded structure of **tp** (Fig. 2(c)). Moreover, the correlations between fourth and fifth 0.05 s yield considerably high accuracy on both datasets **tp** (Fig. 7(b) inset). These time periods overlap with the time we expect the brain is processing word and picture stimuli[35].

The power intensity of frequencies between 4 and 13 Hz yields the highest accuracy on both SEN and FST data (Fig. 7(c)), the peak is 6 Hz for SEN and 8 Hz for FST. These peaks roughly corresponds to the Theta and Alpha frequency band which are related to the resting state, memory, and mental coordination [36]. The accuracy is also moderately high on part of Beta band (14-31 Hz) where attention and concentration are recruited [36]. We also grouped the frequencies into canonical frequency bands and discovered a similar pattern (Fig. S11).

### Identifiability changes with data size and preprocessing

The last dimensions that we investigate is the dependence of individual identification on the amount of available data and on the level of data preprocessing.

We look at the identification accuracy using the three brainprints while increasing the sample size *n*. The identification accuracy increases with the amount of data used for computing **sp, fp**, and **fq** (Fig. 8(a)) as the sampling variance becomes smaller. In general, with 50 s of data, the brainprints perform well on cross-session identification of the same task. **sp** becomes reasonably accurate on the HCP dataset with 100 trials corresponding to 250 s of recording, possibly because more trials are required to accurately compute features that are distinguishable within a larger pool of individuals. For FST and SEN, the identification accuracy of **sp** saturates at fewer number of trials than **tp** and **fq**. It is possible that **sp** requires fewer trials to be estimated robustly.

**Figure 8:**
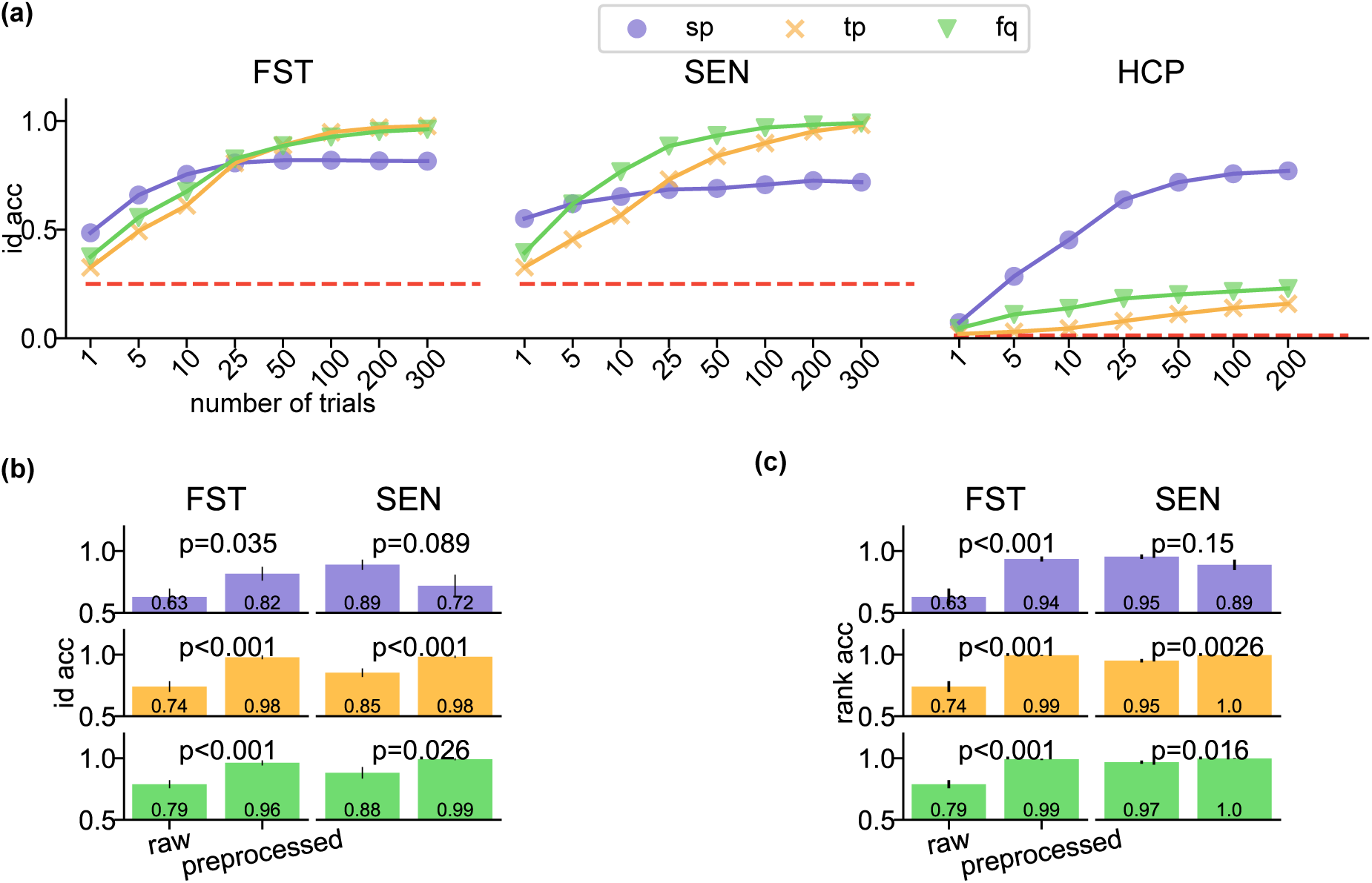
Factors affecting identification accuracy. **(a)** Identification accuracy with respect to the number of trials (sample size) used for the featurization of FST, SEN, and HCP data. Each dot represents the identification accuracy averaged across individuals, identification runs, and cross-sessions (or cross-task sessions) excluding the within-session or within-task results. Error bars are the SEs across the corresponding cross-sessions (or cross-task sessions), individuals, and identification runs of each dataset and are invisible due to small values. **(b)-(c)** Identification **(b)** and rank **(c)** accuracy of the three features computed on raw and fully preprocessed FST and SEN data. The same color represents the same feature as in **(a)**. For **(b)**, the identification accuracy across sessions (*N* = 12 for FST and = 6 for SEN) and individuals (*N* = 4) were averaged with respect to identification runs (*N* = 100) and were put into one vector (of *N* = 48 entries for FST and 24 entries for SEN) for each feature and level of preprocessing. The heights of the bar plots are the mean of the corresponding vector. A two-sided unpaired *t*-test was performed on the vectors of the same feature and dataset between the raw and preprocessed data. The p-values for all pairs are less than 0.05, except for the **sp** feature for SEN. For **(c)**, the rank accuracy were put into one vector in the same way as in **(b)**. The heights of the bar plots are the mean of the corresponding vector and the error bars are its SE A two-sided unpaired *t*-test was performed on the vectors of the same feature and dataset between the raw and preprocessed data. The p-values for all pairs are less than 0.05, except for the **sp** feature for SEN

Preprocessing may also affect identification accuracy. We compared the difference in the identification and rank accuracy between the raw and preprocessed data (Fig. 8 (b,c)). The changes in accuracy were all statistically significant (Fig. 8(b,c))) when the raw data was preprocessed for all the three features except for the **sp** feature for SEN. For both FST and SEN, preprocessing yielded better accuracy for **tp** and **fq**. However, for **sp**, the results point in opposite directions: preprocessing increases identifiability for FST and decreases it for SEN (though with little statistical significance). There was one difference in the preprocessing pipeline for both datasets: FST preprocessing did not include head position correction due to a lack of head position recordings whereas SEN does. Head position correction might be changing the signal in inhomogeneous ways thereby undermining the identifiability with **sp**. We also found that head size and average movement have a weak correlation with identification accuracy in the HCP data, (not statistically significant after multiple comparison correction), shown in Fig. S12.

## Discussion

An individual can be identified with a number of differential characteristics, including their real ‘fingerprints’. Existing studies have suggested the existence a fingerprint in brain signals (e.g. [14, 15]). In this paper, we argued that such brainprints also exist in MEG data and, in fact, there are multiple of them that capture different information from the MEG data. We showed that these brainprints are likely not by-products of environmental artifacts and may pertain to the underlying brain response to stimuli. These analyses, apart from adding to the existing evidence of the brainprints, may bear alarming meanings in privacy issues and provoke thoughts on how scientific conclusions based on multiple individuals have to be examined carefully given these consistent individual-specific features.

In this section, we first discuss the implications of these results in detail, following the same order of the previous section. We then mention limitations and potential improvement to our analysis of brainprints.

### Within-session identifiability

Using the HP data, we showed that both random forest classification with topomaps and 1NN classification with certain interpretable features can be used to correctly identify individuals when the data is collected on a single session. The high accuracy based on merely 0.5 s of data for **sp** and 25 s for **tp** and **fq** is striking since small amounts of data usually leads to inaccurate estimates of these features, unless the underlying patterns are strong. The easy task of identifying individuals on single-session dataset points to strong individual-specific patterns which may or may not be brain-activity related.

### Uniqueness of brainprints

The three features we proposed may not be the only characteristics of MEG data that can be used for individual identification. However, these features represent fundamental aspects of MEG data (and even time series in general) hence they may be a vital first step to understand brainprints. Specifically, we propose the temporal feature, **tp**, because of the high temporal-resolution of MEG data. This feature may have not been used for other types of neuroimaging datasets, suggesting that different features may be informative depending on the nature of dataset of interest.

### Cross-session identifiability

The high cross-session identification accuracy using **sp** confirms it is a brainprint, and supports the previous literature on the similar features in fMRI and EEG [15, 38]. The higher accuracy by **sp, tp** and **fq** suggest that multiple aspects of the individual activity captured by MEG may be used for identification. The generally lower accuracy from **sp** might be the result of the change of alignment of sensors for each individual. However, since necessary steps have taken in the preprocessing pipeline to align the sensors (Supplement A) and each MEG sensor measures brain activity from a non-trivially large area, it remains unclear if the issue is the alignment. Another interpretation of this result is that the temporal and frequency information is more consistent for an individual across time and the spatial information may slowly evolve over time (e.g. when the individual slowly moves during the recording).

Some source-target session pairs have lower identification accuracy than others for **sp** (Fig. 4(b,e)) and the identification accuracy is not necessarily reciprocal, for example, 0.76 vs 1 (mean, session 3 as source, session 2 as target vs session 2 as source, session 3 as target) in FST. The lower accuracy of **sp** of on specific source-target sessions of specific individuals suggests that the identifiability of **sp** may not be uniform over time and individuals.

The three highly identifiable features on FST and SEN represent an alarming message for experimentalists to consider before releasing MEG data. The existence of brainprints are also examples of certain functions of the MEG data with high cross-individual variability preserved across sessions, which has been widely discussed on various types of neuroimaging data [8, 39, 11, 40, 41]. For example, the high accuracy with **tp** suggests the existence of individual variability in their temporal response to the same stimuli. Understanding brainprints will facilitate the understanding of the underlying anatomical and functional variability between individuals.

### Cross-task identifiability

The consistent performance of **sp** on the HCP data is in line with a previous study on fMRI of overlapping individuals that the spatial connectome is preserved between tasks [15]. The rank accuracy of **tp** and **fq** on HCP data indicates the potential of these two features to be consistent within individuals (Fig. 5(c)) because the majority of individuals still have higher than chance rank accuracy than identification accuracy. The current underperformance of these two features, as expected, is likely due to the different temporal dynamics between the resting and task data. This difference may be eliminated by removing the trial part from the task MEG, focusing on inter-trial intervals or baseline periods, and hence boost the identification accuracy of **tp** and **fq**. More complicated matching method may be proposed to further boost the performance of these two brainprints. The within-task identification accuracy (Fig. 4(a)), on the other hand, is still high for all features. With the large pool of participants, the high accuracy confirms the strong individual-specific information contained in the three features within a certain task.

### Cross-modality identifiability

We observed that both the temporal and frequency features can be used to identify participants across modalities (MEG and EEG) with accuracy much higher than chance. This further supports the interpretation that the temporal and frequency brainprints are capturing idiosyncrasies that are specific to the time-course of how a stimulus is processed (both in amplitude and in frequency), outside possible artifacts induced by MEG and EEG machines.

### Interpretability of brainprints

For the three brainprints, higher accuracy seems to be associated with the components of features with more stimuli-driven activity: the occipital lobe, the time around the stimulus, and frequency bands the with highest power intensities (Fig. S4, S5, S6). Indeed, MEG signal is most sensitive to transient, coordinated firings of many neurons that happen after stimulus onset. This commonality indicates the possibility that higher accuracy is related to event-related signals, which in turn suggests that identifiability might be caused by different individuals responding differently to the stimulus. This dependence on stimulus may explain the low accuracy with **tp** and **fq** on HCP data and also suggest that the identifiability originates from brain-related activities instead of session- and individual-specific artifacts.

However, these accuracy patterns of specific components of a feature could also be explained by a signal-to-noise ratio argument: regions, time-points, or frequencies related to stimulus processing correspond to parts of the underlying brain signal with higher amplitudes (while the ambient noise amplitude is constant). It might be that the increase in signal magnitude make the (spatial, temporal or frequency) activity patterns that are specific to a individual more detectable by increasing their amplitude relative to the ambient noise, even if these patterns are not inherently related to stimulus processing and are just consistent features of a individual’s brain activity.

### Sample size and level of preprocessing

**sp** accuracy tends to saturate with fewer number of trials than the other two features on FST and SEN data but with more trials on HCP data (Fig. 8(a)). This difference is likely due to the difference in the maximum accuracy a feature can attain: in HCP data, **tp** and **fq** has much lower maximum accuracy and will reach the peak with smaller number of trials. In FST and SEN data, the spatial pattern may require fewer trials to estimate accurately, as compared to the temporal and frequency features.

The artifact removal and temporal filtering in the preprocessing pipeline might have prevented session-specific noise from contaminating individual-specific features, resulting in higher accuracy for **tp** and **fq**. The seemingly contradictory accuracy on **sp** does not justify our results: identifiability using **sp** increases after prepossessing when not performing head position correction but decreases when performing it. On the one hand, it is expected that head position correction would improve identifiability by recentering each individual’s data to the same position in each session. On the other hand, head position correction may remove individual-specific information such as the head shape, causing the decrease in the accuracy of **sp**. Future work and analysis of additional datasets are required to investigate this result. The difference in the accuracy between raw and preprocessed data suggests, for example, encrypting the data with session-specific noise may lower identification accuracy.

### Limitations

Due to the limited availability of multi-session MEG data, more experiments are needed to generalize our findings to a larger population and other tasks. For example, the cross-session identifiability results depend on 4 subjects and may suffer from high variance. A larger population (with multiple sessions per participant) may benefit the interpretation of brainprints and eventually attribute the high identifiability of certain features to the underlying brain mechanism.

Throughout the paper, we assumed both the target and source datasets had the same pool of participants in the scope of this paper. If we don’t know if one individual from the target set is included in the source set, other classification methods which allow for abstaining from classification (e.g. [42]) may be used to account for the case when no label in the source set can be assigned to the individual. This situation is an example of a more realistic identification problem because an individual’s participation in multiple MEG studies is usually unknown to the public.

### Future solutions

More complicated features can be proposed which combine the spatial, temporal, and frequency information to improve identifiability. For example, functional connectivity at different frequency bands has been used to identify twins from other participants [17]. New feature similarity function that focuses on the structure of the correlation matrices may also be used to improve accuracy [43]. Metric learning [44] can also be used to learn the similarity function in a supervised manner which may boost performance with sufficient labeled MEG data.

On the other hand, given the high identification accuracy with brainprints in this study, privacy-preserving algorithms need to be proposed to account for this privacy issue. Federated learning [45] may be a promising framework as data collected from multiple sessions and sites can be analyzed together without revealing critical information of each specific dataset.

## Methods

### Within- vs cross- session

We called a pair of *source* and *target* sets “within-session” if, for each individual, both datasets were collected in the same visit to the scanner. For example, two blocks of a resting-state recording of a participant collected on the same day are within-session. If the two datasets are collected on different days for each individual, they are “cross-session”. For example, a resting state recording on day 1 and another resting-state recording on day 2 are cross-session. Individuals with within-session data may be easier to identify since the source and target data were collected under almost the same environment.

### Within-session data

The Harry Potter dataset was recorded using the 306-channel whole-head MEG system (Elekta Neuromag, Helsinki, Finland) at the Brain Mapping Center at the University of Pittsburgh. Individuals were asked to read a chapter of Harry Potter [23] while each word was presented for 0.5 s on a screen. There were 5176 trials (words) for each one of the eight individuals. There were 306 sensors at 102 locations where each location has one magnetometer and two planar gradiometers whose signal was averaged. The sampling frequency of the data was 1000 Hz which was further downsampled to 200 Hz. Details about the preprocessing of all the datasets in this paper can be found in Supplement A. The data was parsed into trials where each trial corresponds to the MEG recording when an individual was reading a word. Specifically, the trials of individual *k* is 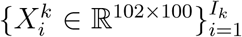 where *I*_*k*_ is the number of trials for individual *k*, 102 represents the number of spatial channels, and 100 represents the number of temporal points in the trial. Since the recording of each individual was collected in one session, we simply split the data into a target and source dataset for the within-session identification task.

### Cross-session data

We considered the following two datasets which have recordings on multiple days:

2- FST data [28], shared online:^1^ individuals saw faces with each face appearing on the screen. There were 1464 trials for each individual. Each trial lasted 0.5 s.

There were 4 individuals and 4 sessions. The sampling frequency was 1000 Hz and was downsampled to 200 Hz. Intervals between consecutive sessions were several days.

2- SEN data (unpublished anonymized citation): individuals read sentences. Each trial lasted 0.5 s. There were 4 individuals and 3 sessions. There were 3575 trials for each individual. The sampling frequency was 1000 Hz and was downsampled to 200 Hz. Intervals between consecutive sessions ranged from days to weeks. In this dataset, two sessions for two individuals were recorded at the same day.

The 306-channel whole-head MEG system (Elekta Neuromag, Helsinki, Finland) at the Brain Mapping Center at the University of Pittsburgh was used to obtain both recordings. The shape of one trial of the two datasets is 102 channels by 100 time points, the same as the Harry Potter data. We used 300 trials to create features for each run of identification. For the within-session identification (diagonal entries of Fig. 3(b,c)), we split the recording for each individual into non-overlapping source and target set before featurization.

### Task vs resting data

We looked at the Human Connectome Project data^2^[5]. The recording was obtained using the whole head MAGNES 3600 (4D Neuroimaging, San Diego, CA) system located at the Saint Louis University (SLU) medical campus during a single-day visit for each individual. There were two sessions, one resting-state recording and one working-memory (WM) task recording where the stimuli were images for the participants to remember. The number of trials for each individual varied because different trials were removed for each individual in the preprocessing step due to signal quality. There were 422 trials in average for the resting dataset and 372 trials in average for the WM dataset. Each trial of the WM corresponded to the 2.5 s of the recording after the onset of the stimulus. The two datasets had 77 individuals in common and we only looked at these individuals. There were 146 channels (after bad-channel removal) and the signal was downsampled to 200 Hz. The two sessions were collected on the same day with a break of several hours. We used 200 trials for featurization for each run of identification due to fewer number of total trials as compared to the aforementioned datasets.

### MEG vs EEG data

We looked at the scene viewing data ^3^. The dataset includes both MEG and EEG recordings of 14 participants viewing 362 scene images (trials), and the MEG and EEG sessions were recorded separately on different days. MEG data was recorded using the 306-channel whole-head MEG system (Elekta Neuromag, Helsinki, Finland) at the Brain Mapping Center at the University of Pittsburgh. EEG data was recorded using a 128-channel whole-head system (ActiveTwo, Biosemi, Amsterdam, Netherlands) at the EEG laboratory of the Psychology Department at Carnegie Mellon University. Each image was shown for 3 − 6 repetitions and the signal was averaged across these repetitions for each image. MEG data was recorded at 1000 Hz and was downsampled to 110 Hz. EEG data was recorded at 512 Hz and was downsampled to 110 Hz. Each trial corresponds to the 1 second of the stimulus presentation. The shape of one trial is 102 channels by 110 time points for MEG (averaged across three sensors for the 102 sensor locations) and 128 channels by 110 for EEG. We used 200 trials to create features for each of the 100 identification runs. For the within-session identification (diagonal entries of Fig. 7(a)), we split the recording for each individual into non-overlapping source and target set before featurization.

### Random forest identification with raw features

We trained a random forest classifier with 256 estimators by first concatenating all the trials of each individual along the time dimension, resulting in **X**_*i*_ ∈ ℝ^102×*N*^, *i* = 1, ⋯, 8 where *N* = 100×5176 is the total number of time points for each individual. We then randomly selected 10000 topomaps from each individual and obtained {**x**_*ij*_ ∈ ℝ^102^, *i* = 1, ⋯, 8, *j* = 1, ⋯, 10000} as the training dataset where each sample is a flattened vector with 102 entries corresponding to the signal across all channels at one time point (or, the topomap). The training label is *y*_*ij*_ = *i*. A 10-fold cross validation was used to compute the classification accuracy. Data was z-scored by channel separately on training and testing data to avoid data leakage. To further avoid dependency between time points, we repeated the analysis while only sampling one topomap per each trial: for each individual, we randomly sampled 5000 trials without replacement and randomly sampled one topomap from each one of the 5000 trials, leading to 5000 randomly sampled topomaps for each individual.

### Interpretable MEG features

Let *X* ∈ ℝ^102×100×*n*^ represent the recording used for featurization, with 102 channels, 100 time points, and *n* randomly sampled trials. The three features are defined as follows:

1. Spatial correlation (**sp**): Pearson correlation between channels averaged over time. *X* was reshaped into ℝ^102×100*n*^ before the correlations between rows of the reshaped matrix were computed.
2. Temporal correlation (**tp**): Pearson correlation between time points averaged over channels. *X* was reshaped into ℝ^100×102*n*^ before the correlations between rows of the reshaped matrix were computed.
3. Frequency (**fq**): power spectrum averaged over channels. Power spectrum of *X*(*i*, :, *j*) was computed using a Tukey window with shape parameter of 0.25 and window size of 100 time points for *i* = 1, ⋯, 102, *j* = 1, ⋯, *n*. The final power spectrum was obtained by averaging across *i, j*.

### Identification using 1NN

We performed *R* = 100 identification runs. In identification run *r*, we randomly split the Harry Potter dataset into non-overlapping source and target set, z-scored the source and target by channel separately, and computed the feature 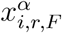 averaged over *n* = 300 randomly sampled trials using data *α* ∈ {target, source} for individual *i* and *F* ∈ {**sp, tp, fq**}. The features from the target to the source set were matched with a labeling with replacement protocol :

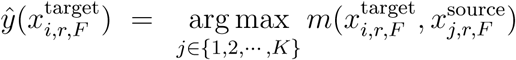

where *K* = 8 is the total number of individuals and *m*(·, ·) is the similarity function measuring the similarity between the two features. We used Pearson correlation as our similarity function. The identification accuracy for individual *i* and feature *F* is 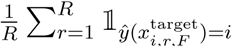 The averaged identification accuracy for feature *F* is 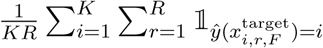. The random baseline is 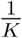.

When the source set and target set were from the same session, we split the dataset into non-overlapping sets as we did in the within-session identification. We did not split data when the source and target data are from different sessions since there is no potential data leakage. We z-scored the data by channel on the source and target separately.

### Rank accuracy

The rank accuracy of individual *i* on one run of identification (suppressing notations of feature *F* and run *r*) is defined as 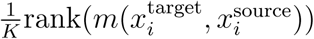 where *K* is the number of individuals, rank 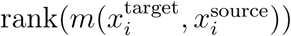 is over 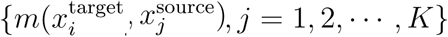. The rank accuracy equals to 1 if the feature of the same individual has the largest similarity between the source and target sets among all *K* individuals, and is 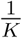 if the similarity is the smallest. The rank accuracy captures more information in a failure case where an individual is misidentified. The random baseline for the rank accuracy is 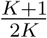.

### Differential identifiability

We also adopted a type of accuracy called differential identifiability [29] to better understand the robustness of the identification. For one identification run, let *C* ∈ ℝ^*K*×*K*^ denote the correlation matrix between the source and target features where *K* is the number of individuals. The differential identifiability for this identification run was calculated as:

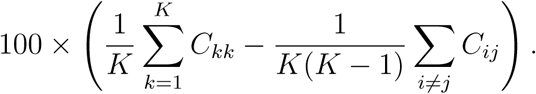

A high differential identifiability would indicate that the features for the same individual (diagonal entries of *C*) are much more similar as compared to features between different individuals (off-diagonal entries of *C*). The chance differential identifiability is 0 where there is no difference between the diagonal and off-diagonal entries if *C*.

### Sub-features

Each feature was decomposed as follows:

1. **sp**: The sensors were partitioned into 8 subgroups according to the map in Figure 1 of [46]: Left Frontal (LF), Right Frontal (RF), Left Temporal (LT), Right Temporal (RT), Left Parietal (LP), Right Parietal (RP), Left Occipital (LO), Right Occipital (LO). Each subfeature was the rows and columns of the spatial correlation matrix corresponding to the sensors in one of the eight groups: letΣ_*s*_ ∈ ℝ^102×102^ be the spatial correlation matrix, then the subfeature corresponding to the correlation between RT and LT, for example, isΣ_*s*_(*ind*_*RT*_, *ind*_*LT*_) where *ind*_*RT*_ is the set of channel indices in the RT group and *ind*_*LT*_ corresponds to the LT group.
2. **tp**: The 100 temporal points were divided into 10 consecutive segments containing 10 time points. Each subfeature was the rows and columns of the temporal correlation matrix corresponding to one of the ten segments: letΣ_*t*_ ∈ ℝ^100×100^ be the spatial correlation matrix, then subfeature corresponding to the correlation between the first and second time segment, for example, isΣ_*t*_(1 : 10, 11 : 20).
3. **fq**: Each subfeature was the segment of the frequency feature vector corresponding to [*f, f* + 10] Hz where *f* ∈ {0, 2,⋯, 90} Hz.

### Raw vs preprocessed data

In Fig. 8(b), we compared the identification accuracy between the raw and preprocessed data for FST and SEN dataset. The details of the full preprocessing pipeline is included in the supplement. In FST dataset, for a given feature and dataset, there were 48 matching results (12 cross-session comparisons ×4 individuals averaged across 100 identification runs) where each one corresponds to the result of deciding which individual from the source session matches the one individual from the target session. An two-sided unpaired t-test was performed to determine if there is a significant difference in the identification accuracy between the raw and preprocessed data. In SEN dataset, for a given feature and dataset, there were 24 matching results (6 cross-session comparisons ×4 individuals averaged across 100 identification runs) instead. The analysis was performed in a similar way for the rank accuracy in In Fig. 8(c).

## Supplementary Material

### A. Data preprocessing

Here we list the preprocessing steps applied to the four types of datasets: Harry Potter (HP), SEN, FST, and Human Connectome Project (HCP). For the EEG and MEG data, please refer to this website for the details of preprocessing: https://figshare.com/articles/dataset/MEG_EEG_data_viewing_scene_pictures/16766938. A summary is listed in Table S1. For all datasets, we used an order 8 Chebyshev type I anti-aliasing filter in Python Scipy package [47] for downsampling. For any within-session identification task, data was z-scored within its corresponding type of dataset (target vs source). Some steps of preprocessing were performed using the python MNE package [37].

1. **HP/SEN**: The 306-channel Elekta Neuromag system was used for the recording. Source-space separation (SSS) along with Maxwell filtering and their temporal extension (tSSS) [48, 49] were used for bad channel correction, head position correction, and electromagnetic artifacts removal. Empty room artifacts were removed. 1 ∼ 150 Hz bandpass filter and 60 & 120 Hz notch filter were used to remove line noise. Heartbeats and eyeblinks artifacts were removed with signal-space projection (SSP) [50]. The data was downsampled to 200 Hz and z-scored by channel within each individual and session.
2. **FST** (preprocessing pipeline was included in the source code): The 306-channel Elekta Neuromag system was used for the recording. Source-space separation (SSS) along with Maxwell filtering and their temporal extension (tSSS) were used for bad channel correction and electromagnetic artifacts removal. Empty room artifacts were removed. We didn’t perform head position correction since there was no head position data. 1 ∼ 150 Hz Bandpass filter and 60 & 120 Hz Notch filter were used to remove line noise. Heartbeats and eyeblinks artifacts were also removed with SSP. The data was downsampled to 200 Hz and z-scored by channel within each individual and session.
3. **HCP**: Both resting and WM datasets were already preprocessed and downloaded from the HCP database^4^. The details of the preprocessing pipeline can be found at https://www.humanconnectome.org/storage/app/media/documentation/s1200/HCP_S1200_Release_Reference_Manual.pdf. MAGNES 3600 (4D Neuroimaging, San Diego, CA) system was used for the recording. For WM data, we looked at the TIM partition which corresponds to −1.5 ∼ 2.5 s relative to the onset of the image. For both resting and WM data, the sampling frequency of the preprocessed data is 508.63 Hz, and 2 s of data were selected from each trial. This corresponds to the whole 1018 time points in the resting data and [763 : 1780]-th time point for the WM data (corresponding to 0 ∼ 2 s relative to the onset of the image). The 2 s data was then downsampled to 101.73 Hz. Data was z-scored by channel within each individual and each data type (resting and WM). We looked at the 146 channels which were marked “good” among all the 77 overlapping individual between resting and WM.

**Table S1:** Summary of the preprocessing stpes for HP, SEN, FST, and HCP data

| Steps | HP/SEN | FST | HCP |
| --- | --- | --- | --- |
| bad data | corrected | corrected | removed |
| head position | corrected | not corrected | not corrected<sup>5</sup> |
| electromagnetic artifacts | removed using SSS | removed using SSS | removed with bad data |
| empty room artifacts | removed | removed | removed<sup>6</sup> |
| band filtering | 1 ∼ 150 Hz | 1 ∼ 150 Hz | 1.3 ∼ 150 Hz |
| notch filtering | 60 & 120 Hz | 60 & 120 Hz | 59 – 61&119 – 121 Hz |
| ECG (heartbeat) artifacts | removed with SSP | removed with SSP | removed with ICA |
| EOG (eyeblink) artifacts | removed with SSP | removed with SSP | removed with ICA |
| downsampling | 200 Hz | 200 Hz | 101.73 Hz |
| z-scoring | by channel within individual, session | same | same |
| shape of a trial<br>[channels, timepoints] | [102, 100] | [102, 100] | [146, 204] |

<sup>5</sup>No continuous recording of head position was available in HCP data
<sup>6</sup>page 68 of https://www.humanconnectome.org/storage/app/media/documentation/s1200/HCP_S1200_Release_Reference_Manual.pdf

### B. Statistical significance of the results

The identification and rank accuracy were averaged across subjects, identification runs, and session pairs. The reported accuracies, being so large, are both statistically and practically significant, and are nearly impossible to attribute to random chance, but accurately quantifying the uncertainty is challenging in our setup. Since featurization for each session of each subject was done before the matching, there is some weak dependence on the accuracy between subjects, session, and identification runs. This dependence makes it hard to analytically obtain a p-value for the accuracy. One numerical alternative is to permute the original recording within each session across subjects before performing matching, but this is computationally expensive as it involves loading and computing large chunks of data 1000s of times. Hence we provide below a (natural, but approximate) permutation-based method for a p-value to test the null that the match is a random guess.

Let **y**^*i*^ denote the true labels of session *i*. Note that **y**^*i*^ = [1, 2, 3, 4]^*T*^ for any session.

The permutation test is performed as follows:

#### Algorithm 1: Null distribution for the identification/rank accuracy

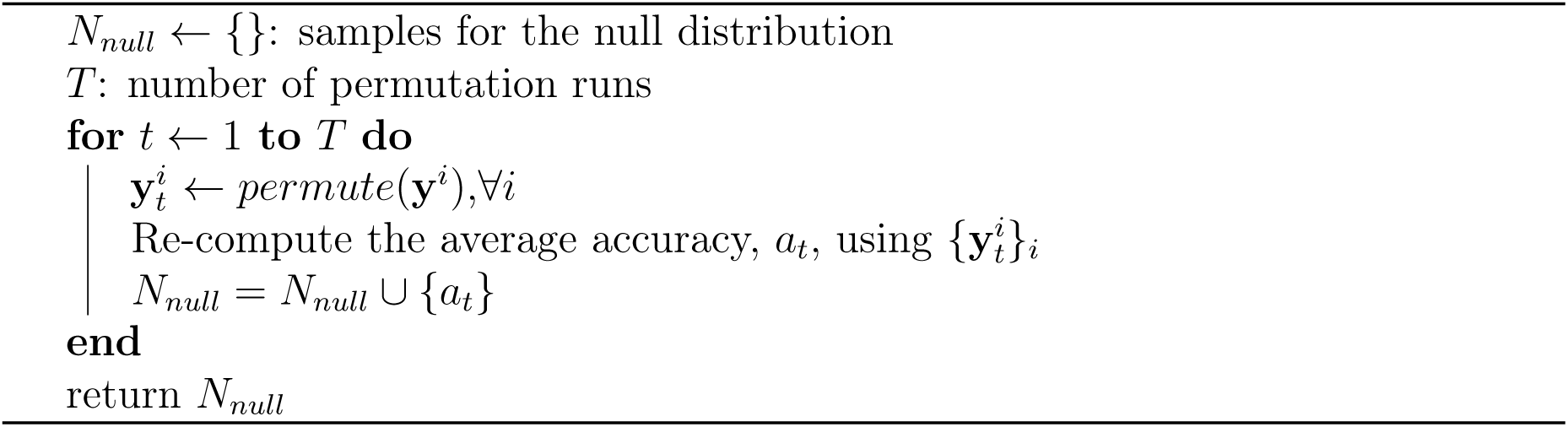

To calculate the p-value, we simply compute 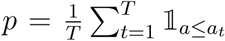, where *a* is the observed average accuracy of a feature across subjects, sessions, and identification runs. Algorithm 1 permutes the labels for each session independently but the permutation remains unchanged for the same source-target pair across identification runs. We summarize the p-values for the identification and rank accuracy of three features on the FST, SEN, and HCP data using *T* = 4999 permutation runs. For all the p-values, since we have not encountered any *a*_*t*_ that exceeds the accuracy number, their values are simply 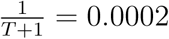. We emphasize that even though these p-values are technically only approximate due to some weak dependence, the fact that we did not see a single permutation which achieved a higher accuracy than ours should convince even rigorous skeptics that it is nearly impossible to explain away our accuracies to chance.

**Table S2:** Statistical significance for the accuracy numbers

| **Data** | **Feature** | **id acc** | **p-val: id** | **rank acc** | **p-val: rank** |
| --- | --- | --- | --- | --- | --- |
| FST | **sp** | 0.816 | 0.0002 | 0.936 | 0.0002 |
| FST | **tp** | 0.978 | 0.0002 | 0.994 | 0.0002 |
| FST | **fq** | 0.962 | 0.0002 | 0.991 | 0.0002 |
| SEN | **sp** | 0.719 | 0.0002 | 0.888 | 0.0002 |
| SEN | **tp** | 0.983 | 0.0002 | 0.996 | 0.0002 |
| SEN | **fq** | 0.991 | 0.0002 | 0.998 | 0.0002 |
| HCP | **sp** | 0.771 | 0.0002 | 0.974 | 0.0002 |
| HCP | **tp** | 0.159 | 0.0002 | 0.819 | 0.0002 |
| HCP | **fq** | 0.229 | 0.0002 | 0.845 | 0.0002 |

## C. Supplementary figures

**Figure S1:**
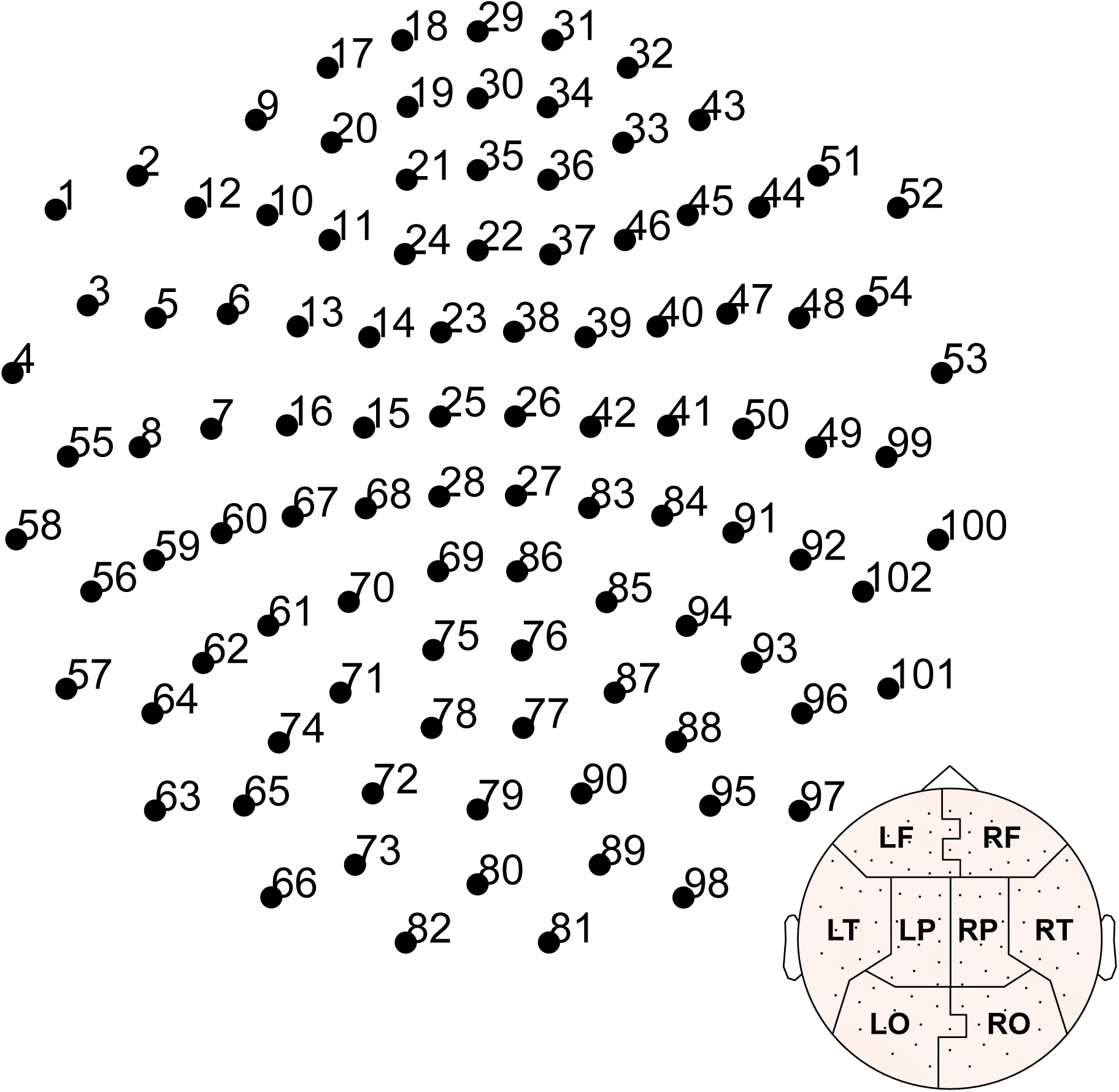
Layout of the sensors for HP,FST, SEN, and MEG/EEG(MEG) data (306-channel Elekta Neuromag system). The channel numbers are consistent with the channel id of **sp** (if specified). Inset is the partitioning of the sensors same as Fig. 7(a) of the main text.

**Figure S2:**
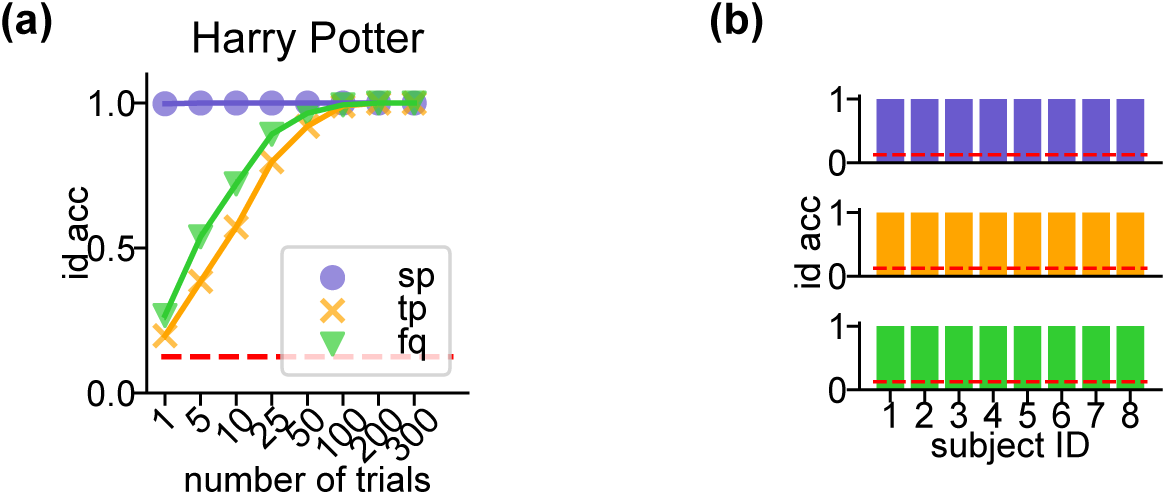
Identification accuracy of sp,tp, and fq on the Harry Potter data. **(a)** Related to Fig. 3. Each dot was averaged across individuals (= 8) and identification runs (= 100). Error bars are the SEs across individuals and identification runs and are invisible due to small values. Each trial is 0.5 s in length. The trends for **tp** and **fq** are similar to that of the cross-session data (SEN and FST). **sp** requires as few as one trial to achieve a perfect accuracy. This indicates strong spatial patterns in the HP data which are specific to each individual. This is expected since HP does not have more than one session, and the identification accuracy for **sp** may be lower if there are multiple sessions in HP data, similar to what we have observed on FST and SEN data. **(b)** Identification accuracy per individual for the three features on HP data.

**Figure S3:**
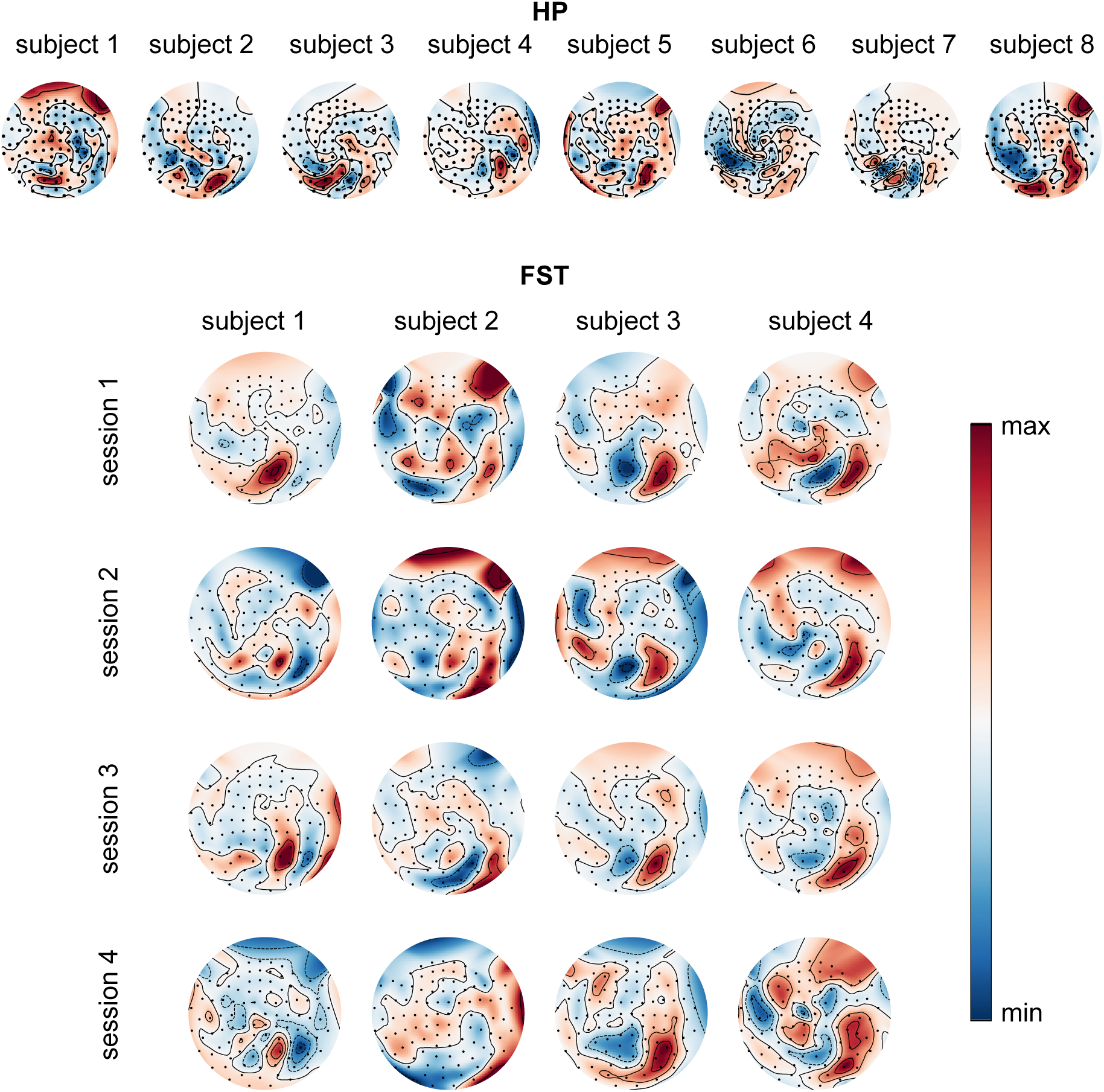
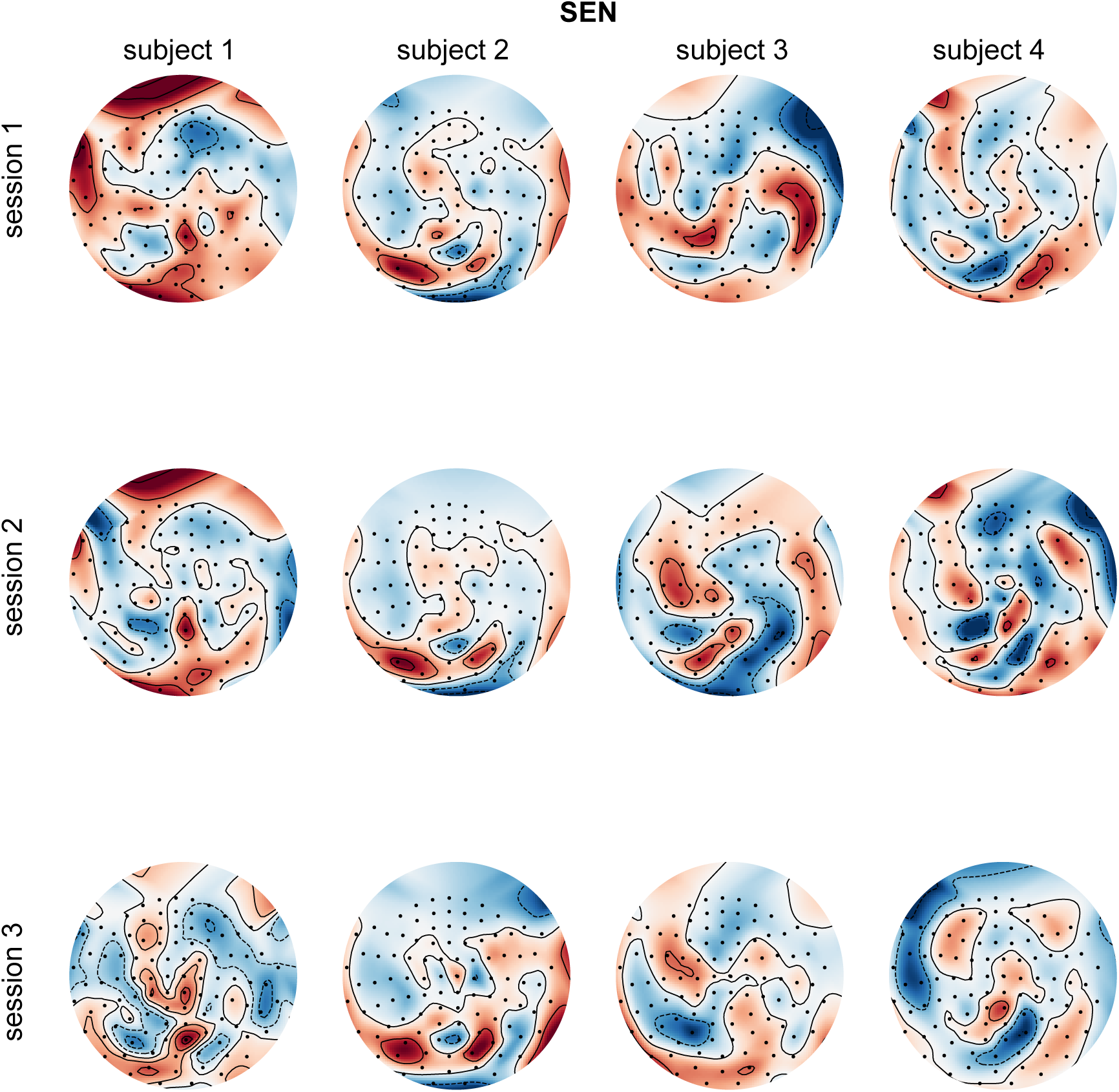
Example topomaps of HP, FST, and SEN data. We plotted the topomap of these three datasets because they were all recorded using the 306-channel whole-head MEG system (Elekta Neuromag, Helsinki, Finland). Each topomap represents the spatial distribution of the brain activity at *t* = 100 ms into a trial, averaged across all trials in a session for each subject. The minimum and maximum of the colormap were specific to each topomap and for convenience, we use one colorbar to represent the scale of the color. For the multi-session FST and SEN data, the spatial distribution of the signal of an individual is more similar across sessions (within a column) as compared to that of other individuals (between columns).

**Figure S4:**
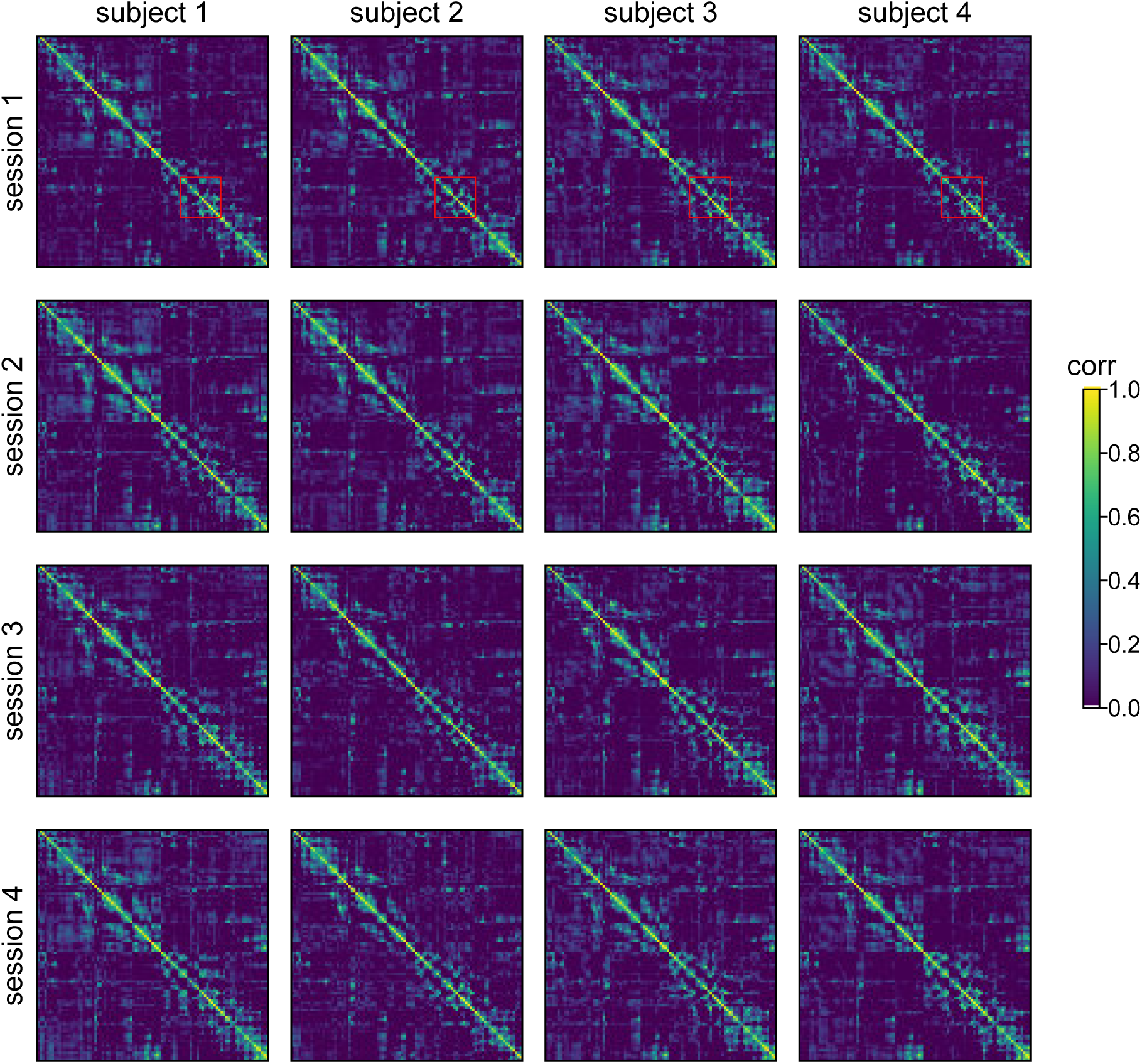
Example sp (spatial connectivity) of FST data. Each heatmap represents a 102 × 102 spatial correlation matrix, where each entry represents the correlation between the brain activity on the corresponding channels. For better illustration we clipped the correlation into [0,1]. The general patterns of the correlation matrices are similar to each other. Some subsets of the heatmap, for example, the bottom-right corner, the top-left corner, and the red rectangle areas are more consistent within a individual and different between individuals. This suggests that only the interactions among a subset of sensors are individual-specific. The red rectangle areas, in particular, roughly correspond to the correlations within the left occipital (LO) lobe which yields the highest identification accuracy on both FST and SEN data (see Fig. 7(a) and Fig. S10). More complicated comparison algorithms may be proposed to focus on these specific subsets to improve the identification accuracy.

**Figure S5:**
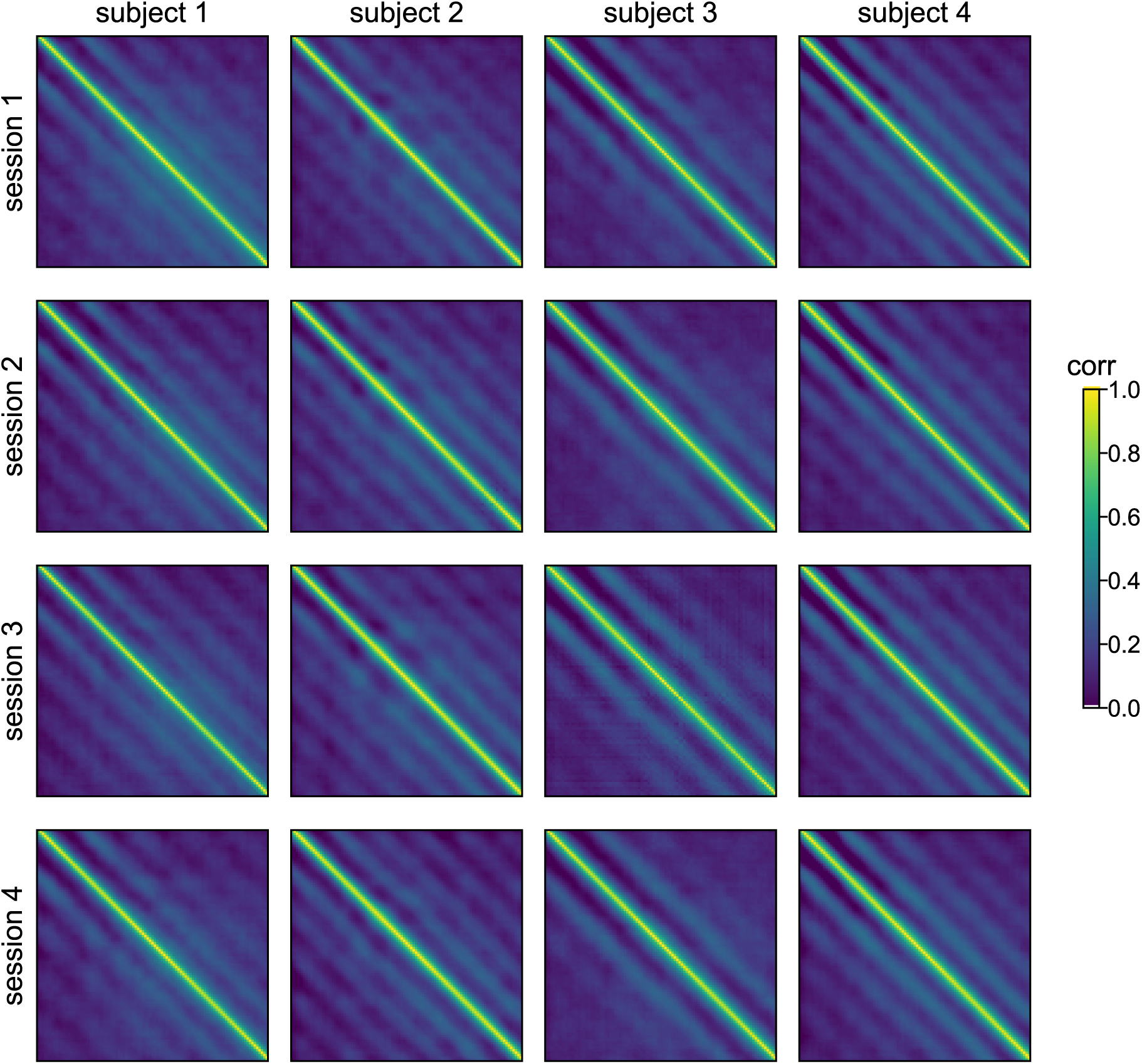
Example tp (temporal connectivity) of FST data. Each heatmap represents a 100 × 100 temporal correlation matrix, where each entry represents the correlation between the brain activity on the corresponding time points. For better illustration we clipped the correlation into [0,1]. The banded structure of the matrices are preserved for the same individual across sessions, and are different between individuals in terms of the number of bands and the relative locations of the bands. The banded structure indicates that there are stronger correlations of the signal with itself at certain lags. In other words, looking at the auto-correlation of the signal or even cross-correlation between different channels may reveal interesting results about the temporal dynamics of the brain activities. The individual-specific band structures also confirm the findings in Fig. 7(b) that correlations of the signal with itself at certain lags are best able to identify individuals.

**Figure S6:**
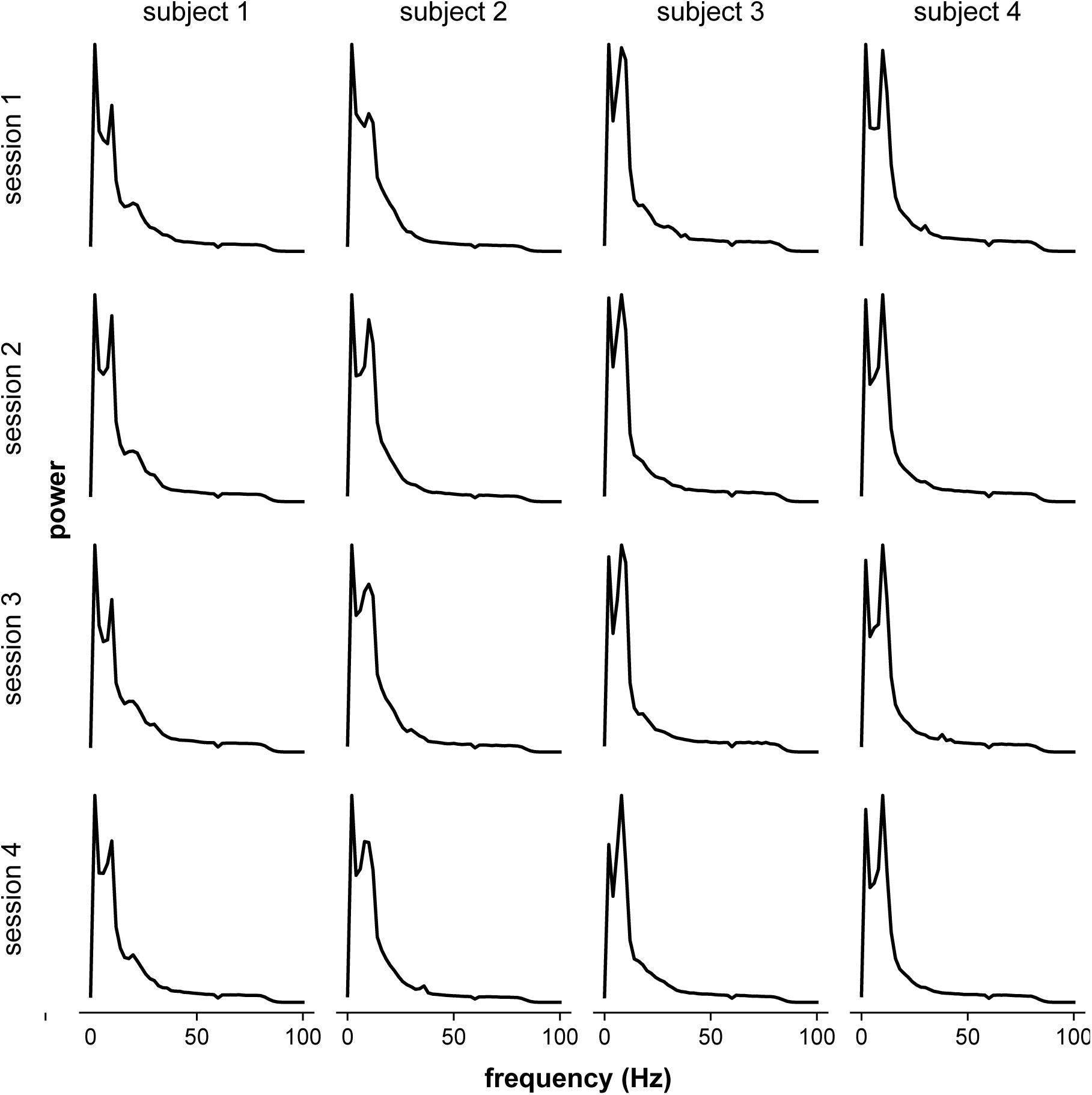
Example fq (frequency) of FST data. Each plot represents power (averaged across channels) vs. frequencies (Hz), where the range of frequencies is [0, 100] with a 2 Hz increment. For all individuals, there are two peaks in the power spectrum. The two peaks correspond to around 5 and 10 Hz. The relative height of the two peaks as well as the shape of the curve near the two peaks are consistently unique to an individual across sessions and different across individuals. There are also small peaks near 20 Hz for some individuals. These frequencies with higher amplitudes seem to align with the results shown in Fig. 7(c) where the frequency band near 10 Hz yields the highest identification accuracy. Hence the components of **fq** associated with more stimuli-driven activity or larger signal-to-noise ratio seem to yield better results.

**Figure S7:**
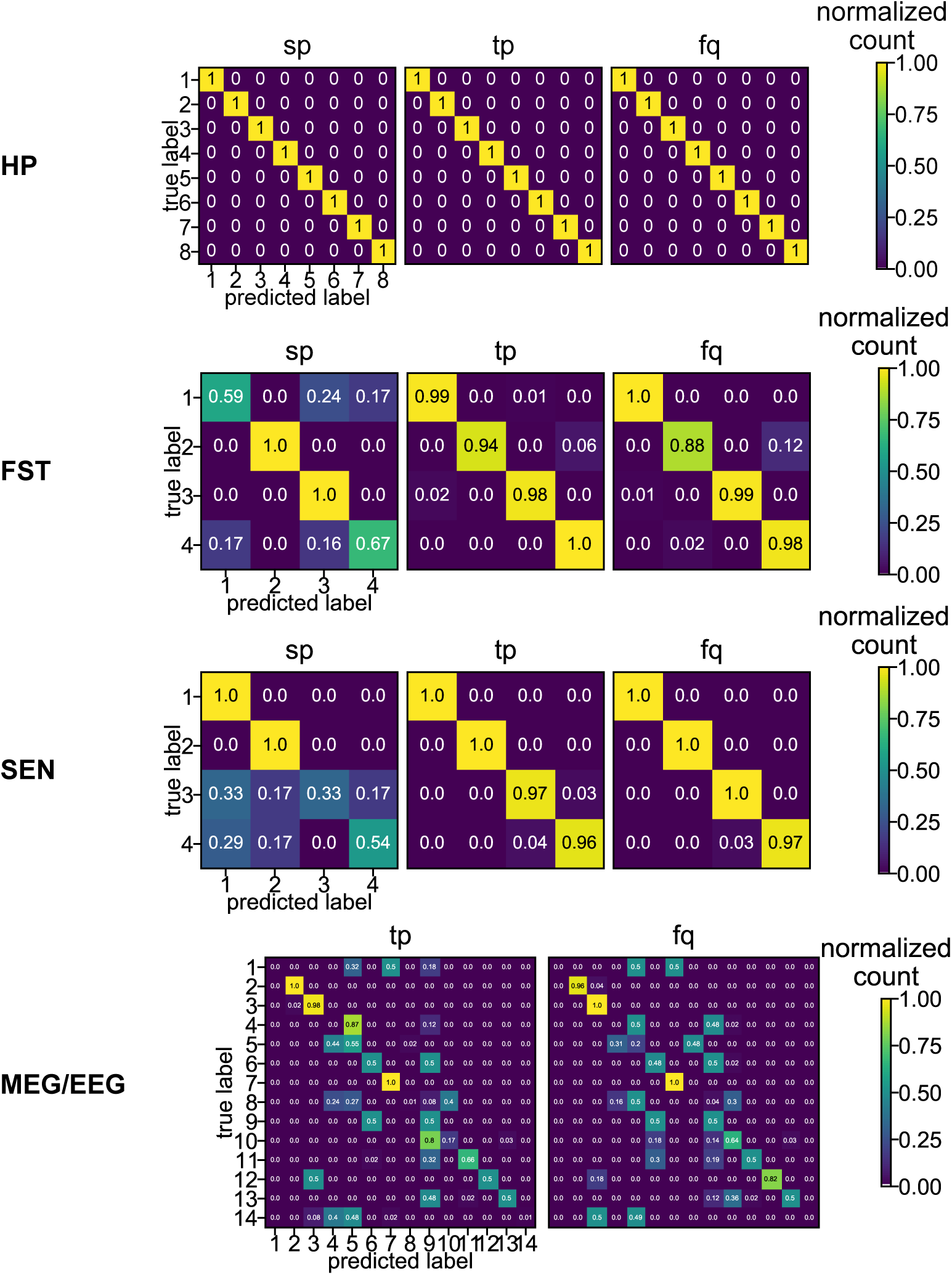
Confusion matrices for HP, FST, SEN, and MEG/EEG data. For the HP, FST, SEN, and MEG/EEG datasets, we constructed the confusion matrices from the 100 identification runs of 2 cross-sessions (HP), 12 cross-sessions (FST), 6 cross-sessions (SEN), and 2 cross-modality (MEG/EEG). The numbers were normalized by the sum of each row and represent the percentage of true labels predicted as the corresponding label on the horizontal axis.

**Figure S8:**
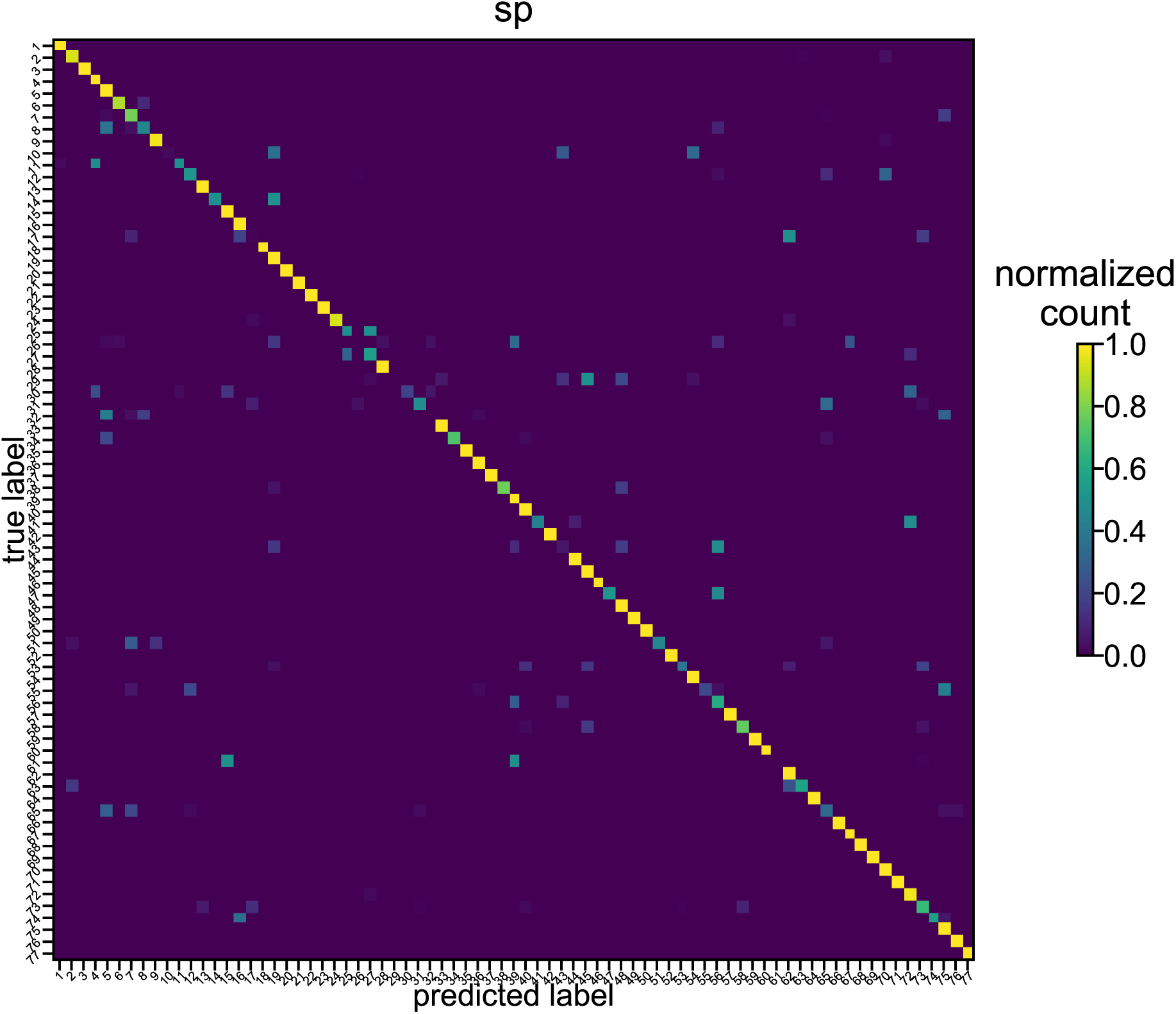
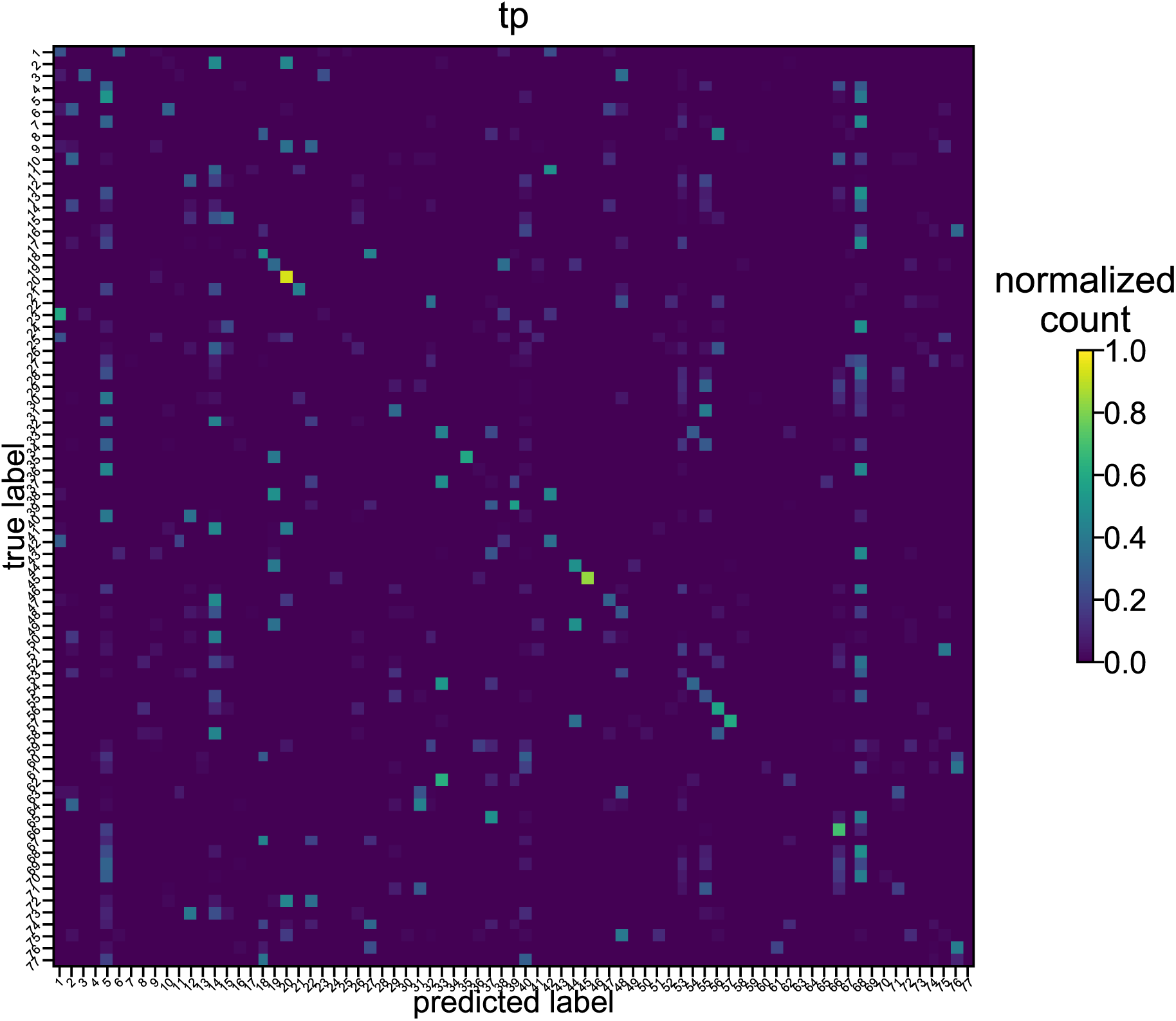
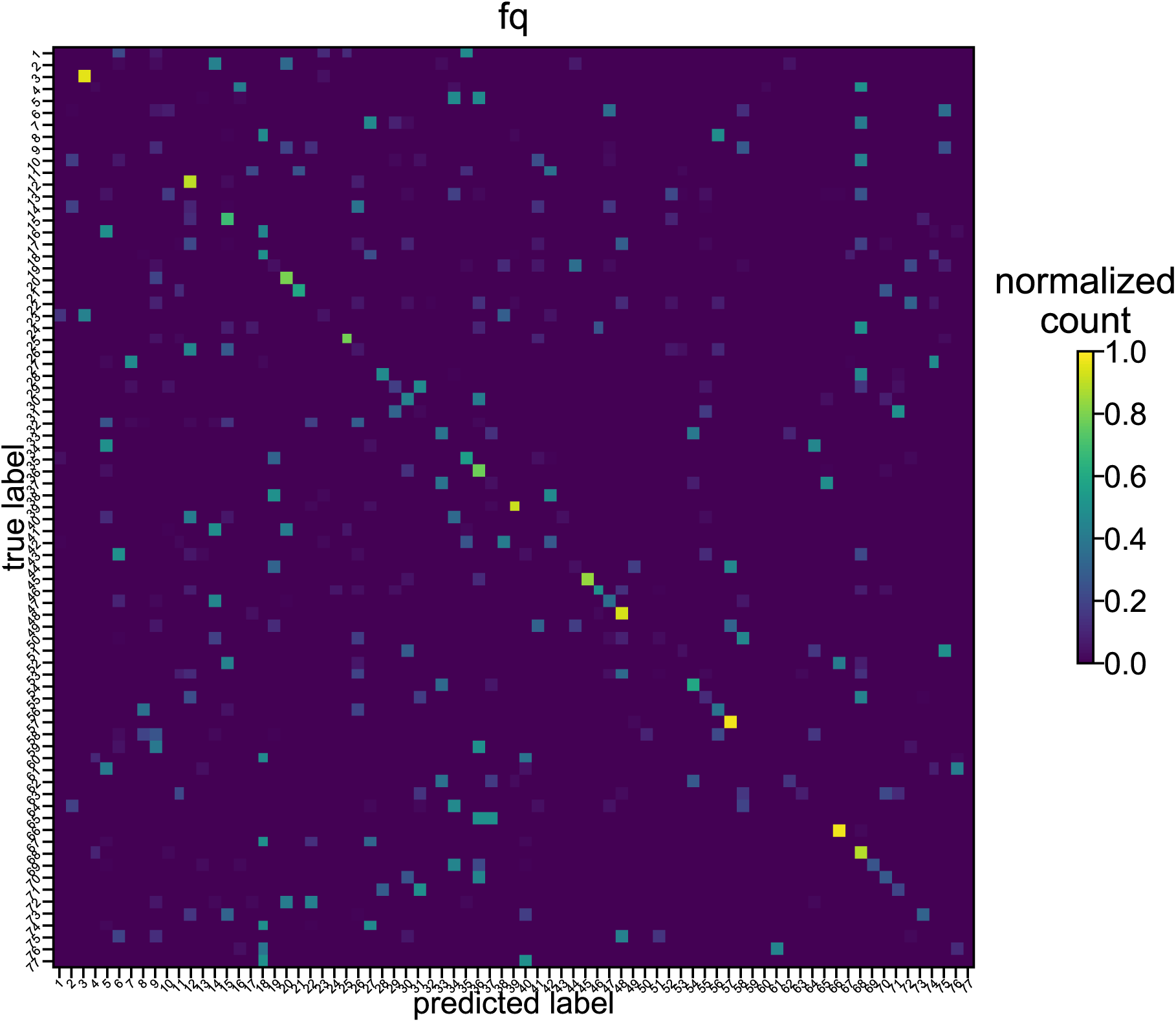
Confusion matrices for HCP data. For the HCP datasets, we constructed the confusion matrices for the three features from the 100 identification runs of 2 cross-sessions. The numbers were normalized by the sum of each row and represent the percentage of true label predicted as the corresponding label on the horizontal axis.

**Figure S9:**
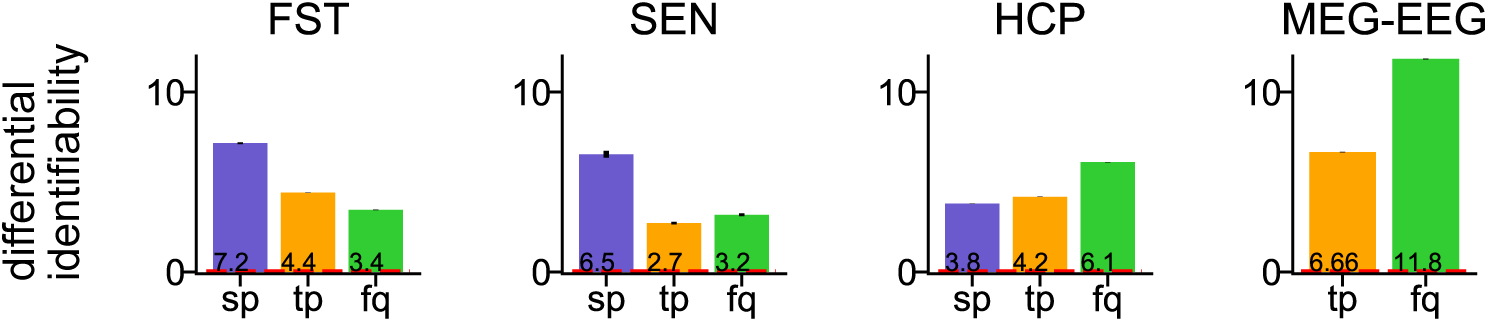
Differential identifiability of the four datasets. We computed the differential identifiability of the FST, SEN, HCP, and MEG-EEG datasets. The random baseline is 0 (red dashed line). The error bar represents the SE across cross-sessions and identification runs and is invisible on some datasets. The differential identifiability for all the features considered is much larger than the random baseline on all the datasets. For FST and SEN, even though **sp** has lower identification accuracy than the other two features, their differential identifiability is still higher. This phenomenon is flipped for the HCP and MEG-EEG datasets. This might be due to the fact that many components of **tp** and **fq** are similar for the same task (i.e. in FST and SEN), leading to smaller deferential identifiability than **sp**.

**Figure S10:**
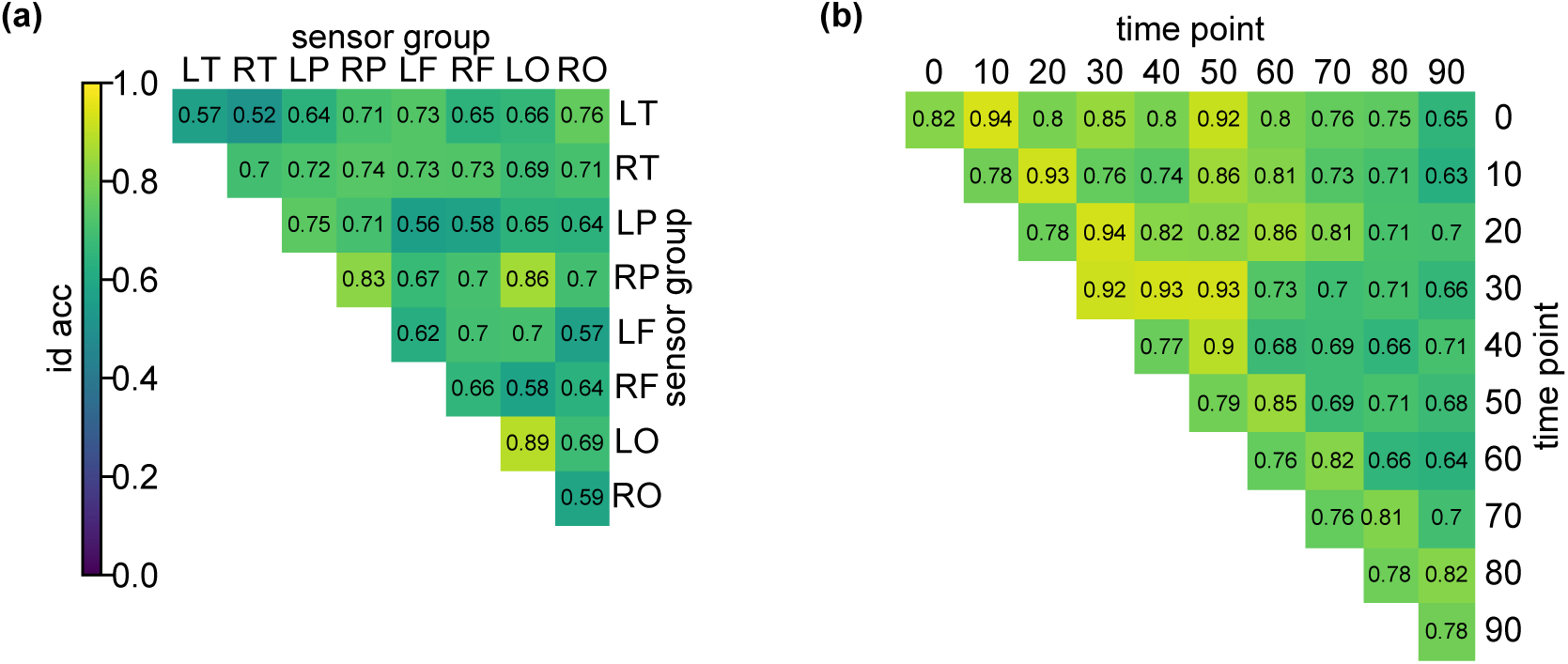
Identification accuracy with sub features for a: sp and b: tp in FST data. Related to Fig. 7. For both FST and SEN, the within-LO and LO-RP correlations yield high identification accuracy. Similarly, for both FST and SEN, the super-diagonal and the correlations between the fourth and fifth 0.05 s yield high accuracy. The consistency of the results on the two datasets suggest that our conclusions regarding the importance of sub-features are not due to experiment-specific artifacts.

**Figure S11:**
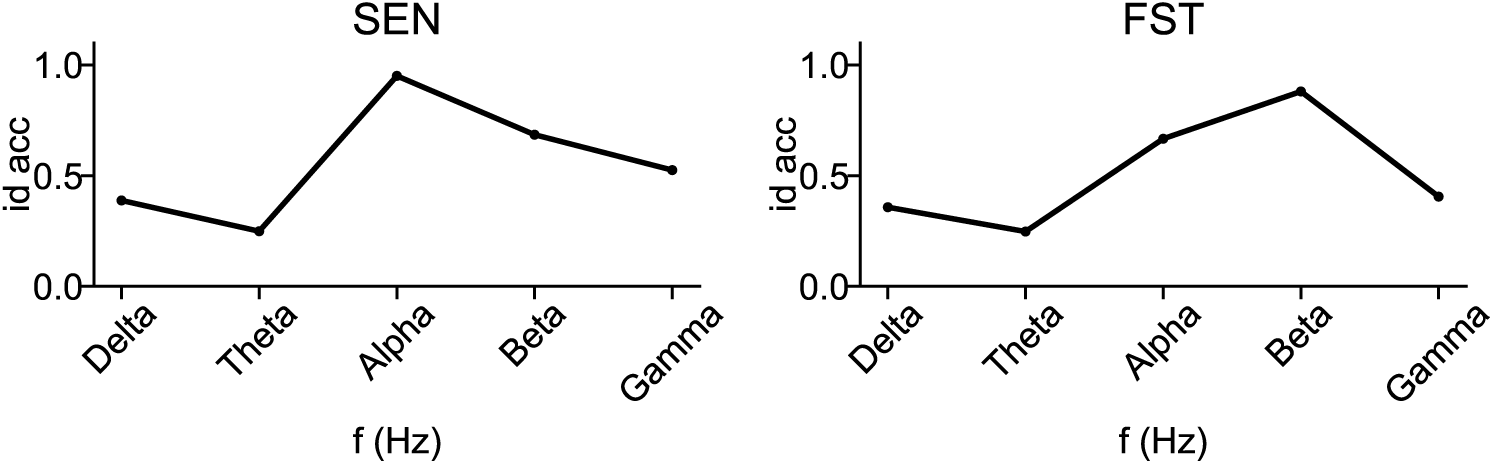
Identification accuracy with respect to frequency bands. Related to Fig. 7. We plotted the identification accuracy with respect to different frequency bands to understand their significance on identifiability. Due to the limitation of sampling frequency, we grouped the frequencies according to this modified range: Delta (0 ∼4 Hz), Theta (4 ∼8 Hz), Alpha (8 ∼14 Hz), Beta (14 ∼32 Hz), and Gamma (32 ∼100 Hz). The different peaks for SEN and FST datasets might be due to the borderline difference between Alpha and Beta bands.

**Figure S12:**
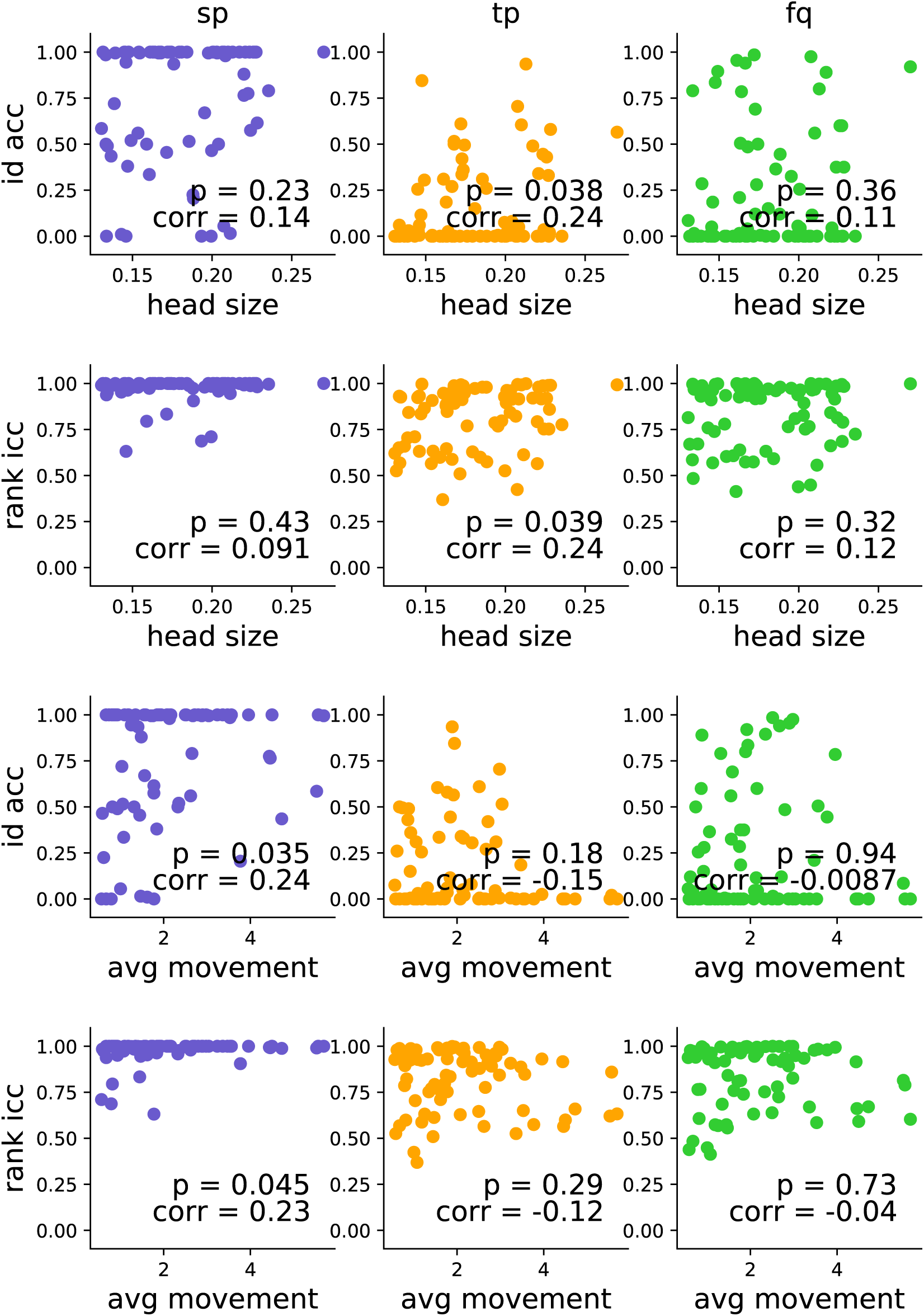
Effect of head size and average movement on identifiability for the HCP data. For the HCP dataset, we looked at how identifiability may change with respect to the size of the individual’s skull and average movement, which were both recorded for each experiment session. We plotted the identification accuracy and rank accuracy with respect to these metrics and computed the statistical significance at the level of individuals (Pearson’s *r*). It can be seen that the average movement may have a positive impact on identification using **sp** possibly due to the fact that more movement may induce more individual-specific information. Bigger head size may lead to higher accuracy using **tp** possibly due to the more distant brain areas recorded and the more accurate average response to the stimuli. The **fq** is not affected by either metric because it is a purely spectral feature and may not be easily influenced by these spatial-related metrics. However, we do ask for caution in interpreting these results because these correlations are not statistically significant after multiple comparison corrections.

## Footnotes

1 https://figshare.com/articles/FST_raw_data/4233107

2 https://www.humanconnectome.org/study/hcp-young-adult

3 https://figshare.com/articles/dataset/MEG_EEG_data_viewing_scene_pictures/16766938

4 https://www.humanconnectome.org/study/hcp-young-adult

5 No continuous recording of head position was available in HCP data

6 page 68 of https://www.humanconnectome.org/storage/app/media/documentation/s1200/HCP_S1200_Release_Reference_Manual.pdf

## Notes

### Competing Interest Statement

The authors have declared no competing interest.

### Summary of Updates

Figures, text, results.

